# Adaptive divergence generates distinct plastic responses in two closely related *Senecio* species

**DOI:** 10.1101/2020.01.24.918201

**Authors:** Greg M. Walter, James Clark, Antonia Cristaudo, Delia Terranova, Bruno Nevado, Stefania Catara, Momchil Paunov, Violeta Velikova, Dmitry Filatov, Salvatore Cozzolino, Simon J. Hiscock, Jon R. Bridle

## Abstract

The evolution of plastic responses to external cues allows species to track the environmental variation they regularly experience. However, it remains unclear how plasticity evolves during adaptation. To test whether distinct patterns of plasticity is associated with recent adaptive divergence, we quantified plasticity for two closely related but ecologically divergent Sicilian daisy species (*Senecio*, Asteraceae). We sampled c.40 genotypes of each species from natural populations on and around Mt Etna and then reciprocally transplanted multiple clones of each genotype into four field sites along an elevational gradient representing each species’ native range, the edge of their range, and conditions outside their native range. At each elevation we quantified survival and measured leaf traits that included investment (specific leaf area), morphology, chlorophyll fluorescence, pigment content and gene expression. As evidence of adaptive divergence, both species performed better at their native site and better than the species from the other habitat. Traits and differentially expressed genes that changed with elevation in one species often showed little change in the other species, or changed in the opposite direction. Adaptive divergence is therefore associated with the evolution of distinct plastic responses to environmental variation, despite these two species sharing a recent common ancestor.

## Introduction

The resilience of many natural populations to novel or changing environments relies on their ability to adjust their phenotype to track local conditions (Chevin et al. 2010). This phenotypic plasticity generates different phenotypes from the same genotype depending on the environment to which it is exposed (Via et al. 1995; Ghalambor et al. 2007; Charmantier et al. 2008). The ability for plasticity to help a species track environmental variation is shaped by selection within environments routinely experienced by that given species (Ghalambor et al. 2007). Plasticity is therefore only likely to buffer populations by maintaining fitness when exposed to familiar environmental regimes (Bradshaw 1965; Schlichting 1986; Baythavong and Stanton 2010), which will likely affect how populations respond to novel environmental conditions (Chevin and Hoffmann 2017). Characterising plasticity for closely related, but ecologically divergent species can test to what extent adaptive divergence is also associated with divergence in phenotypic plasticity.

The effect of adaptive divergence on plasticity will depend on how selection interacts with plasticity (de Jong 2005; Radersma et al. 2020). In particular, phylogenetic relatedness (Pigliucci et al. 1999; Kellermann et al. 2018), ecology (Kulkarni et al. 2011) and the predictability of the environment (Oostra et al. 2018) can determine the nature and amount of variation in plastic responses among given taxa, as well as the range of environments within which plastic responses are adaptive. Although contrasting environments are expected to select for different forms and different magnitudes of plasticity (Schlichting 1986; Donohue et al. 2001), it is not known to what extent a shared common ancestry will constrain such divergence in plasticity. Few studies have assessed whether adaptation to contrasting environments causes not only adaptive divergence, but also divergence in plastic responses when exposed to the same environmental variation.

Assaying variation in gene expression across environments can reveal key aspects of the genomic basis of plasticity. As allelic (sequence changes in regulatory genes) or epiallelic (e.g. DNA methylation, chromatin remodelling, post-transcriptional modifications) variation underlying trait plasticity becomes fixed during adaptation to a particular environment, differences in plasticity are likely to arise among populations occupying contrasting environments (Gibson and Wagner 2000; Shaw et al. 2014). Highly predictable environments will reduce genetic variation in plasticity through stabilising selection acting either on the genetic regulators (e.g. transcription factors) or long-term epiallelic changes, such as transgenerational DNA methylation (Colicchio et al. 2015; Oostra et al. 2018). If such stabilising selection varies among environments, divergence in phenotypic plasticity will occur. Determining the gene expression profiles for closely related but ecologically divergent species can therefore reveal how plasticity evolves during adaptive divergence.

In this study, we first use a common garden experiment to identify physiological differences between two closely related species of *Senecio* that inhabit contrasting elevations on Mt. Etna, Sicily. We then quantify variation in survival and phenotypic plasticity across four field transplant sites that lie within and beyond the elevational range of either species. *Senecio chrysanthemifolius* (**Fig. 1a**) is a short-lived perennial (individuals generally live for less than two years) with highly dissected leaves that occupies disturbed habitats (e.g., vineyards, abandoned land and roadsides) in the foothills of Mt. Etna c. 400−1,000m a.s.l (above sea level) and at similar elevations throughout Sicily. By contrast, *S. aethnensis* (**Fig. 1b**) is a longer-lived perennial with entire, glaucous leaves and is endemic to lava flows c. 2,000−2,600m a.s.l on Mt. Etna that are covered by snow each winter. These two species are proposed to have diverged recently, possibly c.150,000 years before present, around the same time that the uplift of Mt. Etna created the high elevation environment in which *S. aethnensis* is found (Chapman et al. 2013; Osborne et al. 2013). Their recent shared ancestry is reflected by very low genetic divergence across the genome, despite large differences in habitat, phenotype and life history (Chapman et al. 2016). In particular, rapid divergence in leaf shape and dissection is key feature of speciation in this system (Brennan et al. 2009), as well as in other *Senecio* species (Walter et al. 2018).

**Fig. 1.**
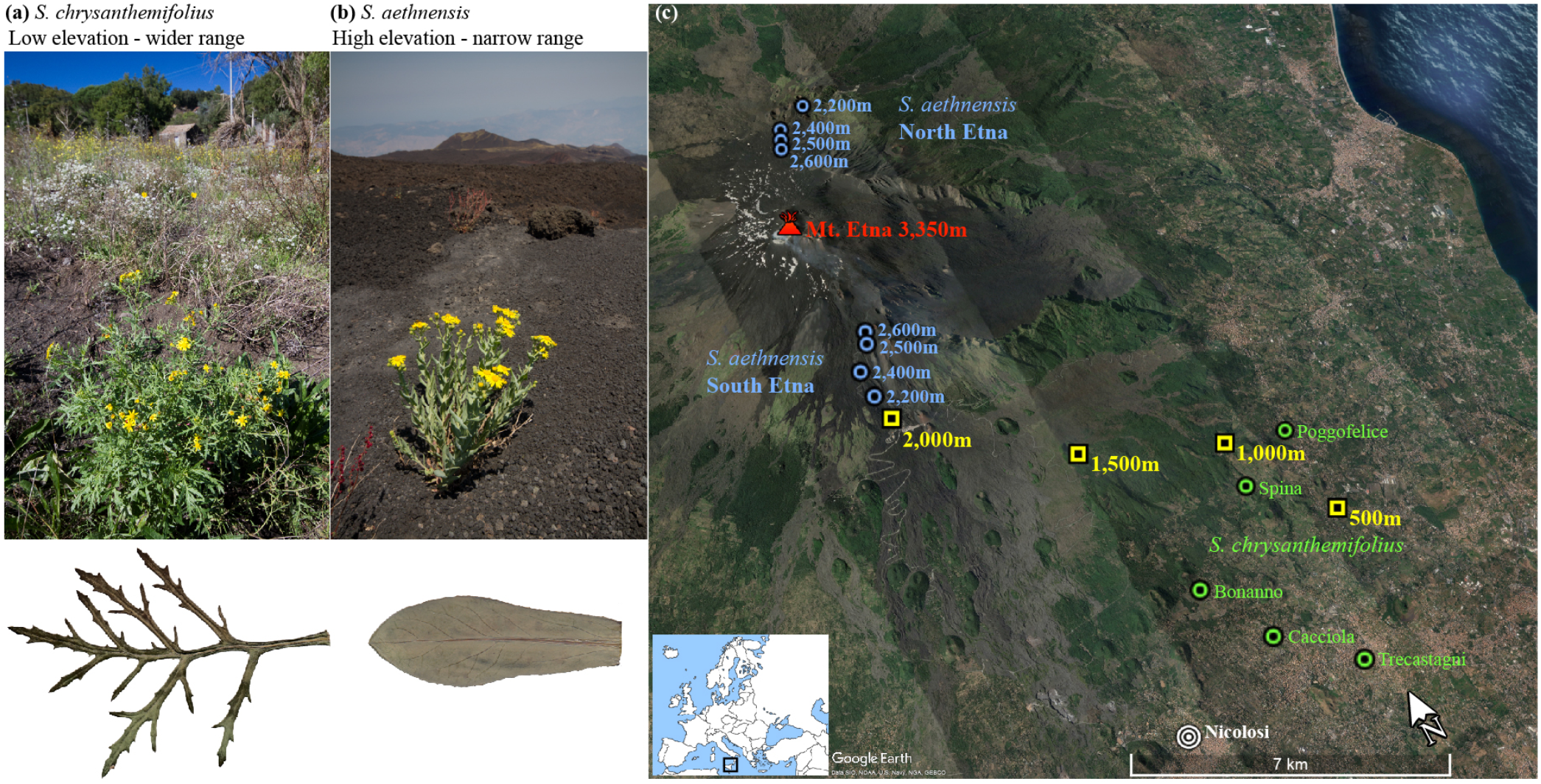
**(a)** *Senecio chrysanthemifolius* occupies disturbed habitats below c.1,000m a.s.l, and has thin, dissected leaves. **(a)** *Senecio aethnensis* inhabits lava flows and has thicker, smooth-margined leaves with a thick waxy cuticle. **(c)** Map of sampling locations (*S. chrysanthemifolius*: green circles; *S. aethnensis*: blue circles) and transplant sites (yellow squares). Sampling locations for *S. aethnensis* and the transplant sites are labelled by their elevation. Inset map shows the location of the study system within Europe.

We predicted that: (1) The occupation of contrasting habitats by these species would be reflected by differences in physiology in a common garden, suggesting functional trait divergence; (2) As evidence of adaptive divergence, species would perform better at their native elevations than beyond their existing range, and better than the other species within their native habitat; (3) Despite their recent common ancestry, and reflecting their adaptive divergence into contrasting habitats, we predicted that the two species would show different patterns of plasticity at both the phenotypic and gene expression level. To test these hypotheses, we sampled 79 genotypes of *S. chrysanthemifolius* and *S. aethnensis* from natural populations and conducted two reciprocal transplant experiments in 2017 and 2019. The relative ease by which both species can be propagated from cuttings means we are able to control for genetic variation by assaying identical genotypes at multiple sites as clones, allowing biological replicates in analyses of both phenotypic traits and gene expression profiles. Both experiments involved transplanting multiple cuttings of each genotype to four transplant sites across an elevational range that included the home range of each species, and two intermediate elevations. Genotypes that were propagated for use in the 2017 experiment were sampled directly from natural populations as cuttings, while genotypes propagated for the 2019 experiment were raised in a common garden from seeds collected from individuals growing in natural populations. In each transplant experiment, we quantified survival, growth, and leaf traits including leaf shape, gene expression, photosynthetic activity, pigment content and leaf investment.

## Materials and methods

### Sampling natural populations

We sampled achenes (i.e. seeds), and took cuttings from naturally growing individuals of both species after plants started flowering. Sampling was conducted in May-June 2017 for *S. chrysanthemifolius* and July 2017 for *S. aethnensis* because *S. aethnensis* grows more slowly and flowers later than *S. chrysanthemifolius*, given its higher elevation habitat. For *S. chrysanthemifolius*, we sampled from 88 individuals from five sites between 500−800 m.a.s.l (**Fig 1c**, **Table S1**). For *S. aethnensis*, we sampled from 87 individuals at four different elevations (2,600m, 2,500m, 2,400m and 2,300m.a.s.l) on both the North and South slopes of Mt. Etna (**Fig 1c**, **Table S1**). Although this species occurs at 2,000m, we avoided sampling below 2,300m to avoid the risk of sampling hybrids associated with a stable hybrid zone present at 1,500−1,700m (Brennan et al. 2009). To minimise the risk of sampling close relatives, most plants sampled were more than 10m apart.

We then used the seeds and cuttings sampled from the individuals in the natural populations for three separate experiments. First, to test whether the two species have evolved differences in physiology under common garden conditions, we germinated seeds and grew plants in the laboratory to measure a suite of physiological traits. Second, to test for differences in plasticity, in 2017 we propagated the cuttings from the field genotypes and reciprocally transplanted them at four elevations on Mt. Etna. Third, to verify the 2017 experiment, we conducted a second reciprocal transplant of cuttings in 2019, but this time we germinated field-collected seeds and grew plants under common garden conditions, from which we propagated cuttings and conducted the transplant.

### Physiological differences between species under common garden conditions

To assess differences in physiology between the species, we grew plants from seeds in a growth cabinet (see **Methods S1**). From eight maternal families of *S. chrysanthemifolius* we grew 34 individuals, and from ten maternal families of *S. aethnensis* we grew 41 individuals. Seeds were scarified mechanically and placed in petri dishes containing moist filter paper. Seedlings were transplanted into 70mm square pots with standard potting mix and grown for two months before physiological measurements were taken. We used a Dualex+**®** instrument (ForceA, France) to measure the leaf content of chlorophyll, anthocyanin and flavonol pigments. Using an LCpro gas analyser (ADC BioScientific, UK), we measured photosynthetic gas exchange. Intrinsic water use efficiency (iWUE) was calculated as the ratio between photosynthesis and stomatal conductance. Chlorophyll fluorescence was measured using an IMAGING-PAM M-series chlorophyll fluorometer (Heinz Walz GmbH, Germany). Using output from the fluorometer, we quantified two mechanisms of physiological light defence of leaves (**Methods S1**): the unregulated [Y(NO)] and regulated [Y(NPQ)] dissipation of heat.

### 2017 field transplant experiment

In the glasshouse (Giarre, Italy), cuttings from all individuals sampled from natural populations (hereafter, genotypes) were cut into 5cm stem segments, each possessing 2-3 leaf nodes. Each smaller cutting was then dipped in rooting plant growth regulator for softwood cuttings (Germon® Bew., Der. NAA 0.5%, L. Gobbi, Italy) and placed in a compressed mix of coconut coir and perlite (1:1) in one cell of an 84-cell tray. All cuttings from each genotype were kept together in one half of a tray and tray positions were randomised regularly. Trays were kept moist and checked regularly for cuttings that successfully produced roots extending out of the bottom of tray. For each genotype, rooted cuttings were randomised into experimental blocks and transplanted at four field sites. From the initial genotypes sampled, we transplanted 37 *S. chrysanthemifolius* genotypes and 42 *S. aethnensis* genotypes.

We transplanted multiple cuttings of each genotype into three experimental blocks at each transplant site. Cuttings were transplanted into grids of 20×7 plants, with the position of cuttings randomised with respect to genotype, and separated from each other by 40cm (*S. chrysanthemifolius* block *n*=109; site *n*=327; total N=1,308; *S. aethnensis* block *n*=130; site *n*=390; total N=1,560). Depending on the number of cuttings that successfully produced roots, we transplanted 6-15 cuttings per genotype at each transplant site (see **Table S1**). The two species were transplanted at different times (*S. chrysanthemifolius* in June-July 2017; *S. aethnensis* in August 2017) because seasonal constraints meant that sampling from natural populations of *S. aethnensis* was only possible a month after *S. chrysanthemifolius* had been sampled. Following each transplant, cuttings were watered daily for three weeks to encourage establishment. To prevent death during high temperatures in July-August (consistently >35°C), we watered cuttings daily during this period, which allowed us to assess the phenotypic responses of genotypes to what were still stressful conditions. We recorded mortality approximately every two weeks and measured the phenotypic traits of all plants at a single time point when both species showed substantial post-transplant growth (November 2017).

In 2019, we repeated the 2017 experiment, but transplanted cuttings of genotypes that were grown from field-collected seed in the glasshouse at the same time for both species, and in the same experimental blocks. This meant that the 2017 experiment reflects differences among adult genotypes sampled directly from natural populations and transplanted at different points in the season. By contrast, the 2019 experiment represents cuttings propagated from genotypes raised from seed in a common garden (glasshouse).

### 2019 field transplant experiment

We germinated five seeds collected from multiple genotypes (*S. aethnensis n=*25; *S. chrysanthemifolius n*=21) that were sampled from the natural populations using the protocol described above. After one week, one seedling from each maternal genotype was chosen at random and placed in 14cm pots containing standard potting media and left to grow under supplemental lighting (25W LED tubes; TSA Technology, Italy). When plants reached c.30cm high, we removed all branches and propagated cuttings for the transplant using the same protocol as the 2017 experiment. Plants were left to re-grow, and then we took a second round of cuttings to increase replication in the transplant experiment.

We prepared the field sites in the same manner as in 2017 and transplanted multiple cuttings of each genotype into four experimental blocks at each elevation. Cuttings were transplanted into grids of 20×7 plants and randomised with respect to species and genotype (i.e. both species were planted together). We transplanted the first round of cuttings 16^th^ April 2019 (3 cuttings/genotype/block; *n=*139 plants/block; N=1,668 plants), and the second round of cuttings 24^th^ May (2 cuttings/genotype/block; *n=*92 plants/block; N=368 plants). Overall, we transplanted 2,036 plants. Initial watering was daily for three weeks, after which it was reduced to once per week to maintain a more natural watering regime than the 2017 experiment.

### Field transplant sites

Field transplant sites were at four elevations (500m, 1,000m, 1,500m and 2,000m a.s.l) along a transect on the south-eastern side of Mt. Etna (**Fig. 1c**). The 500m site was located in a garden among fruit trees, the 1,000m site on an abandoned vineyard among *Quercus ilex*, the 1,500m site among an apple and pear orchard, and the 2,000m site surrounded by pine trees on a lava flow from 1983. Both the native elevations (500m for *S. chrysanthemifolius* and 2,000m for *S. aethnensis*) were located less than 1km from natural populations. Plants of both species were often observed at intermediate elevations, but were never observed within the native range of the other species. There is an elevational transition in soil type from a silty sand at elevations between 500m and 1,500m, to volcanic sand at 2,000m. At each transplant site four data loggers (Tinytag Plus, Gemini Data Loggers, UK) recorded temperature hourly. We took three soil samples at each transplant site, which were analysed for 21 variables that included nutrients, salts and ions (Nucleo Chimico Mediterraneo laboratories, Catania, Italy). To analyse the soil data, we used Multi-Dimensional Scaling (MDS) to calculate the scaled distance between the soil samples taken at all transplant sites.

### Characterising leaf morphology and pigment content

To characterise leaf morphology, we sampled and pressed young but fully expanded leaves from each cutting once they showed extensive growth (at least three months after transplantation). For the 2017 experiment, we sampled 3-5 leaves per cutting in November, and for the 2019 experiment we sampled two leaves in September. Leaves were weighed, and then scanned and morphology quantified using the program Lamina (Bylesjo et al. 2008), which generates estimates of leaf area, perimeter and the number of indentations (leaf serrations and lobes). To estimate the density of indentations along the leaf margin, we standardised the number of indentations by the perimeter. To estimate leaf complexity we calculated perimeter^2^/area, where lower numbers indicate less complex (i.e. more entire) leaves. As measures of leaf investment we included leaf area, and calculated Specific Leaf Area 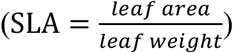, where greater values represent thinner leaves. For the 2019 field experiment, we used a Dualex Scientific+ instrument (Force-A, France) to quantify the concentration of leaf chlorophyll and flavonol pigments. Flavonols are secondary metabolites that help to reduce oxidative stress caused by abiotic (e.g. temperature or light) and biotic (e.g. herbivory) stressors (Mierziak et al. 2014).

### Quantifying chlorophyll fluorescence (2017 field experiment)

To quantify photosynthetic capacity across elevation for both species, we measured chlorophyll *a* fluorescence, which estimates the efficiency of the photosynthetic response to intense light. We selected five genotypes at random from each species and measured chlorophyll fluorescence on four cuttings per genotype at each elevation (site *n=*40 plants, total N=160). We took measurements at two transplant sites each day, completing all four sites within one week in October 2017. For each cutting we measured four leaves, and to temporally replicate measurements we measured the same cuttings at each site on a second day. To take measurements, we put leaf clips on four leaves of each plant and dark-adapted the plants for 30 minutes by covering them with large black plastic containers. We then took fluorescence induction curve measurements for 2 seconds at 3,500μmol s^−1^m^−2^ photosynthetic photon flux density from each leaf (clip) using a Handy PEA instrument (Hansatech Instruments Ltd., UK). We calculated PI_total_, the total performance of photosystem I and II (**Methods S1**).

### Statistical analyses of survival and univariate plasticity

We first tested for significant differences in survival across elevation using mixed effects cox proportional hazards models from the *coxme* v2.2-14 (Therneau 2012) package in R v3.6.1 (R Core Team 2019). Survival at the end of the experiment was used as the response variable (censored at the last point of data collection), and we included transplant elevation and species as fixed effects, and genotype and experimental block as random effects (see equation 1).

We then quantified plasticity as the change in leaf morphology, pigment content and chlorophyll fluorescence across elevation (all leaf traits were first averaged for each clone) using the linear mixed model within the package *lme4* v1.1-23 (Bates et al. 2015)

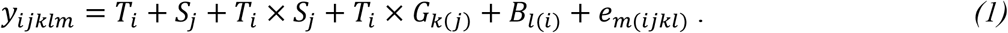

Changes in the response variable across transplant elevation were modelled by the *j*th species (*S_j_*) in the *i*th transplant elevation (*T_i_*) and their interaction (*T_i_* × *S_j_*), which were all included as fixed effects. Random effects included the interaction between transplant elevation and genotype *T_i_* × *G_k(j)_*, which accounted for differences among genotypes at each elevation, and experimental block within each environment (*B_l(i)_*). The residual error variance was captured by *e_m(ijkl)_*. Separate implementations of equation 1 were used for each (normally distributed) univariate response variable of interest (*y_ijklm_*). For each implementation of equation 1, we tested the significance of the interaction between transplant site and species using likelihood ratio tests. To correct for multiple comparisons, we adjusted the P-value by multiplying by the number of tests, and keeping α=0.05. To test whether differences in morphology between transplant sites were significant for each species, we used *emmeans* v1.4 (Lenth 2019) to conduct pairwise t-tests adjusted for multiple comparisons.

### Estimating multivariate plasticity

To quantify multivariate plasticity across elevation, we mean standardised each morphological trait (divided by its mean). We then used a multivariate analysis of variance (MANOVA) to test for significant differences in multivariate mean phenotype across elevation and between species. We analysed each experiment separately by including the morphological traits as a multivariate response variable. We included species, transplant elevation and their interaction as main effects. We included the interaction between transplant site and genotype as the error term, which is the appropriate denominator to calculate F-ratios for the main effects because it tests whether differences among species and transplant elevations are significantly greater than differences among genotypes. To visualise multivariate differences, we used the output of the MANOVA to calculate the D-matrix (**methods S1**), which represents differences in mean multivariate phenotype across species and elevation. We also quantified how the two species differed in their direction of plasticity in response to elevation by calculating the vector of plasticity 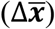 for each species using

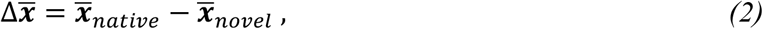

where 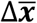 is the vector (standardised to unit length) that represents the change in multivariate phenotype from the native site 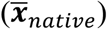 to the novel elevation 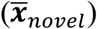, for each species separately (Noble et al. 2019). Calculating the angle that separates 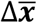 for each species quantifies the difference in the direction of plasticity between species.

### Sampling of plant tissue and RNA extraction

To quantify gene expression variation across elevation and between species, we sampled young leaves from 12 genotypes of each species. For each genotype, we collected 3-4 newly emerged leaves (15-20mm in length) from three cuttings at each transplant site (2 species × 12 genotypes × 3 clones × 4 elevations; total N = 288) in summer (22^nd^ to 26^th^ July 2019), 86-90 days after the initial transplant, which was after cuttings showed sufficient growth. All leaves for a cutting were placed in the same Eppendorf tube and stored in RNAlater at –20°C. We extracted RNA from each sample using QIAgen RNeasy kits. Library preparation and 3’ QuantSeq RNA sequencing was performed at the Oxford Genomics Centre on an Illumina NextSeq platform, producing 75bp single-end reads.

### Quantifying differential expression across transplant sites, genotypes, and species

A reference transcriptome was assembled for each species (**Methods S1**). Trimmed reads were mapped to each species’ reference transcriptome using *Salmon* v0.13.1 (Patro et al. 2017). Read counts were normalised by transcript size, library size and filtered based on counts >5 across half of all samples. All estimates were repeated using *DESeq2* (Love et al. 2014) and *limma/voom* (Law et al. 2014) according to

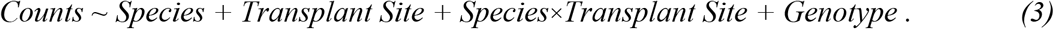

In *limma/voom*, the genotype was modelled as a random effect. For comparisons within species, each treatment was compared with the home transplant site of each species, with differentially expressed genes determined based on an adjusted p-value <0.01 (Benjamini and Hochberg 1995) and a log fold change >2 for overexpression or <−2 for underexpression. In *DESeq2*, log fold changes were shrunk using the ‘apeglm’ method and then used to rank genes based on high overexpression and underexpression.

Reference transcriptomes were annotated (**Methods S1**), and to identify functional categories for differentially expressed genes, gene ontology enrichment analyses were performed using *topGO* v2.3.6 (Alexa and Rahnenführer 2019). Enrichment was determined with a Kolmogarov-Smirnoff test using genes that were significantly differentially expressed (adjusted p values <0.01) between the native elevation and the furthest transplant site.

### Weighted network construction of differentially expressed genes

Weighted Gene Coexpression Network Analysis (WGCNA) identifies correlations of expression among all genes and then forms modules of coexpressed genes within that network. Consensus modules were constructed for each species (**Methods S1**). Each module was then summarized using its first principal component as a module eigengene, which represents the expression profile of the module. We tested for a correlation between each module eigengene in each species and transplant elevation. Each module was tested for gene ontology enrichment in *topGO*, using Fisher’s exact test.

## Results

### Physiological differences between species under common garden conditions

Under common garden conditions, both species show a similar ability to regulate heat dissipation from their leaves [Y(NPQ)] (**Fig. 2a**; t(65)=0.0116, adj. P=1.000), and similar values for non-regulated heat dissipation [Y(NO)] (**Fig. 2b**; t(65)=2.351, adj. P=0.1085). This indicates that photochemical energy conversion and mechanisms protecting leaves from heat are similar for both species under common garden conditions. *Senecio chrysanthemifolius* showed evidence of higher intrinsic water use efficiency than *S. aethnensis* (**Fig. 2b**; t(69)=3.875, adj. P=0.0010), suggesting that leaves of *S. chrysanthemifolius* conserve water more effectively. *Senecio aethnensis* showed slightly greater (but non-significant) leaf concentrations of chlorophyll (**Fig. 2c**; t(143.8)=2.085, adj. P=0.1940), and greater concentrations of flavonols (**Fig. 2d**; t(200.9)=4.399, adj. P<0.0001) than *S. chrysanthemifolius*. Greater concentrations of flavonols indicates a better ability for leaves of *S. aethnensis* to defend against high UV light and lower temperatures (Jugran et al. 2016).

**Fig. 2.**
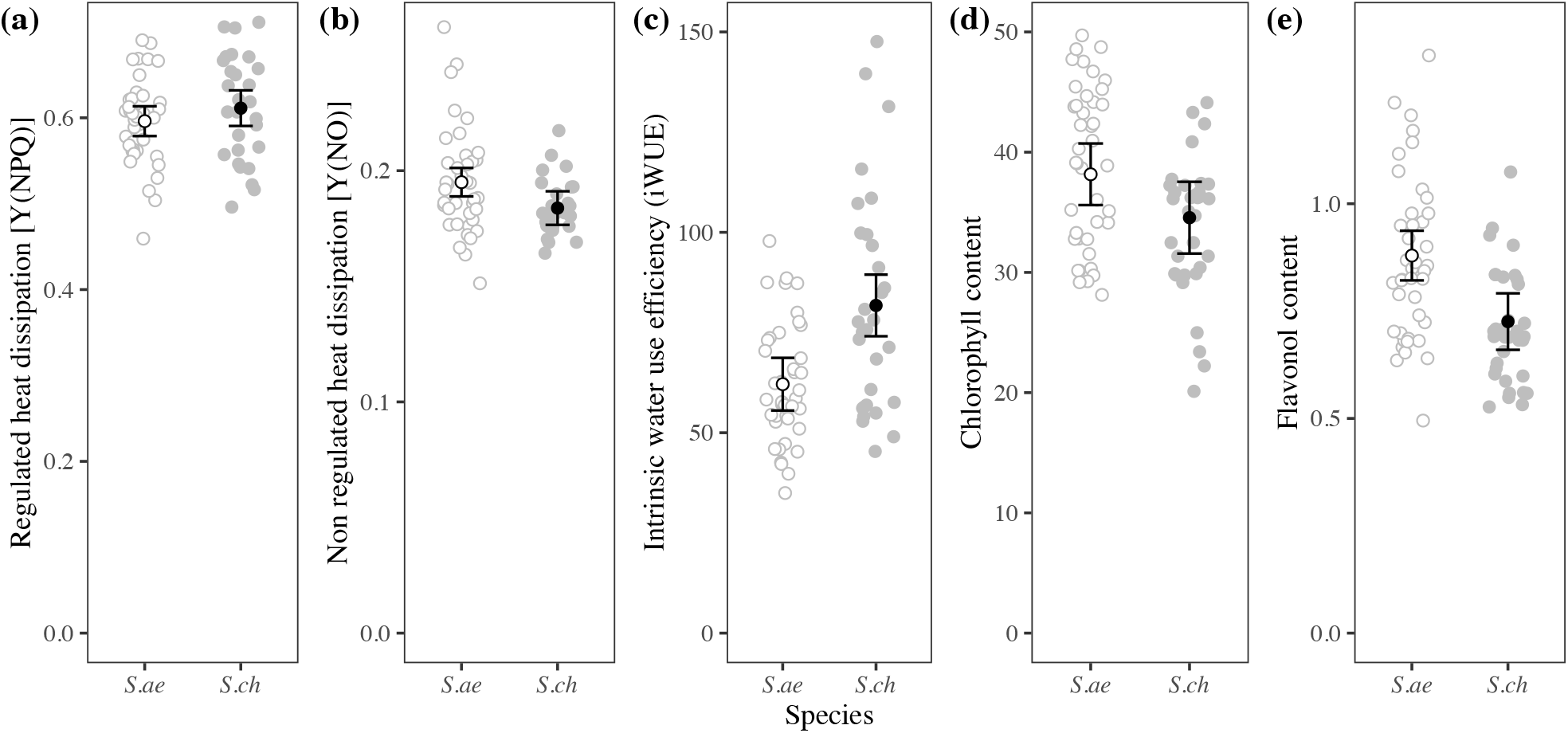
Physiological differences between species grown from seeds under common garden conditions in the laboratory. Filled circles represent *S. chrysanthemifolius* (*S.ch*), while unfilled circles represent *S. aethnensis* (*S.ae*). Gray circles represent individual plants measured and credible intervals represent the 95% confidence intervals of the mean. **(a)** Both species showed similar values for regulated heat dissipation, Y(NPQ). **(b)***Senecio aethnensis* showed greater values for the unregulated dissipation of heat, Y(NO). **(c)***Senecio chrysanthemifolius* showed higher intrinsic water use efficiency, while *S. aethnensis* showed higher leaf chlorophyll content **(d)** and a higher flavonol content **(e)**.

### Survival of transplanted cuttings reflects adaptive divergence between species

The transplant sites experience contrasting climatic conditions associated with elevation, with extreme heat (regularly exceeding 40°C) at 500m and 1,000m during summer, and extreme cold (regularly below 0°C) at 1,500m and 2,000m during winter (**Fig. 3a**). Soil profiles roughly separated the four transplant sites in a linear fashion along the first axis (MDS1), which represented a transition in soil type as a reduction in nutrients (amount of organic material, total nitrogen, cation exchange capacity and exchangeable ions) at higher elevations (**Fig. 3b**; **Table S2**). The second axis (MDS2) characterised differences between the 1,000m site and the other sites, associated with greater concentrations of various salts at 1,000m (soluble nitrates, calcium and magnesium; **Table S2**).

**Fig. 3.**
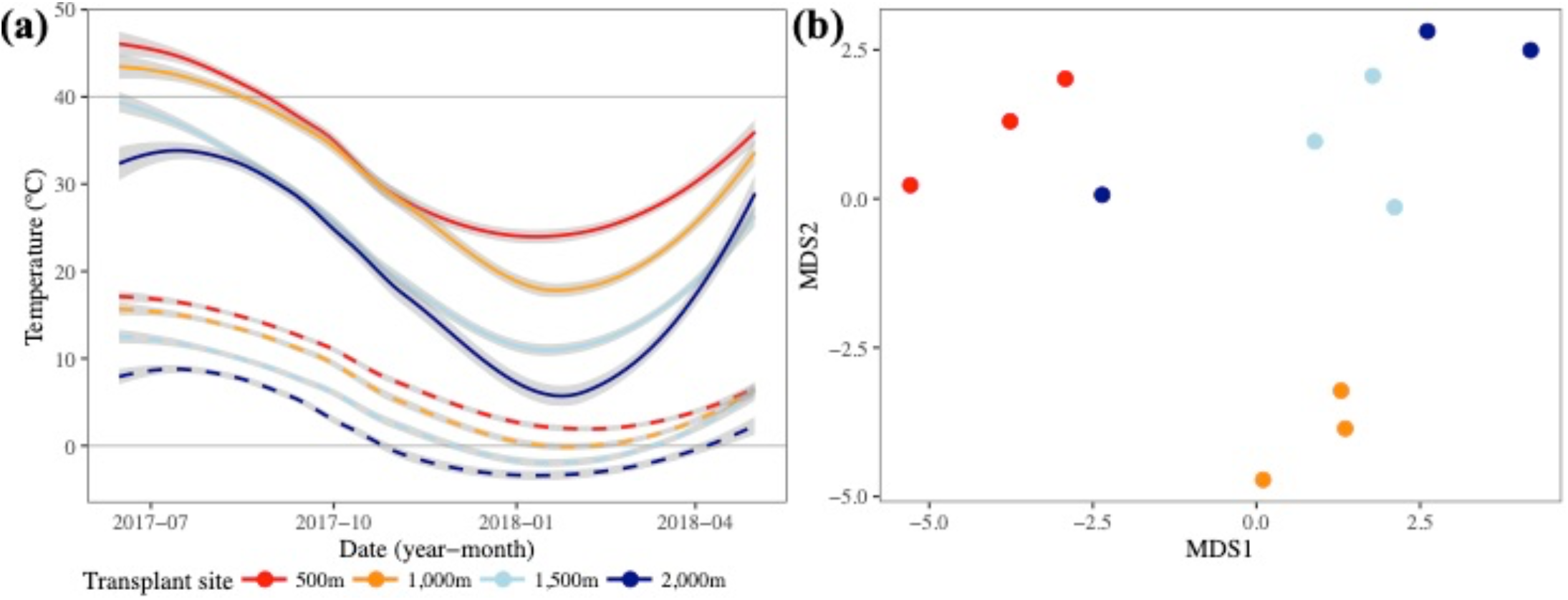
Differences in environment for the four transplant sites at four elevations. **(a)** Average daily maximum (solid lines) and minimum (dashed lines) for three data loggers at each site, for the duration of the transplant. Gray shading represents one standard error in estimating the curves. Higher elevations remained below 40°C in the summer and dropped well below zero in the winter. **(b)** Differences in soil composition for 21 soil variables captured by a multidimensional scaling analysis (**Table S2** presents the soil data).

Both experiments showed strong evidence for adaptive divergence between species whereby each species survived consistently better at their home (‘native’) than the species from the other elevation extreme (**Fig. 4a-b**). Evidence of adaptive divergence was supported by a significant species×elevation interaction for both experiments (2017: *χ^2^*(3)=30.21, P<0.00001, and 2019: *χ^2^*(3)=251.00, P<0.00001). *Senecio chrysanthemifolius* showed significantly greater mortality at higher elevations away from its native 500m site (**Fig. 4a-b**; **Table S3**). By contrast, *S. aethnensis* showed significantly greater mortality at lower elevations away from its native 2,000m site (**Fig. 4a-b**; **Table S3**). In contrast to 2017, in 2019, *S. chrysanthemifolius* suffered c.35% greater mortality at 500m (its native site) than at 1,000m (which is within the native range).

**Fig. 4.**
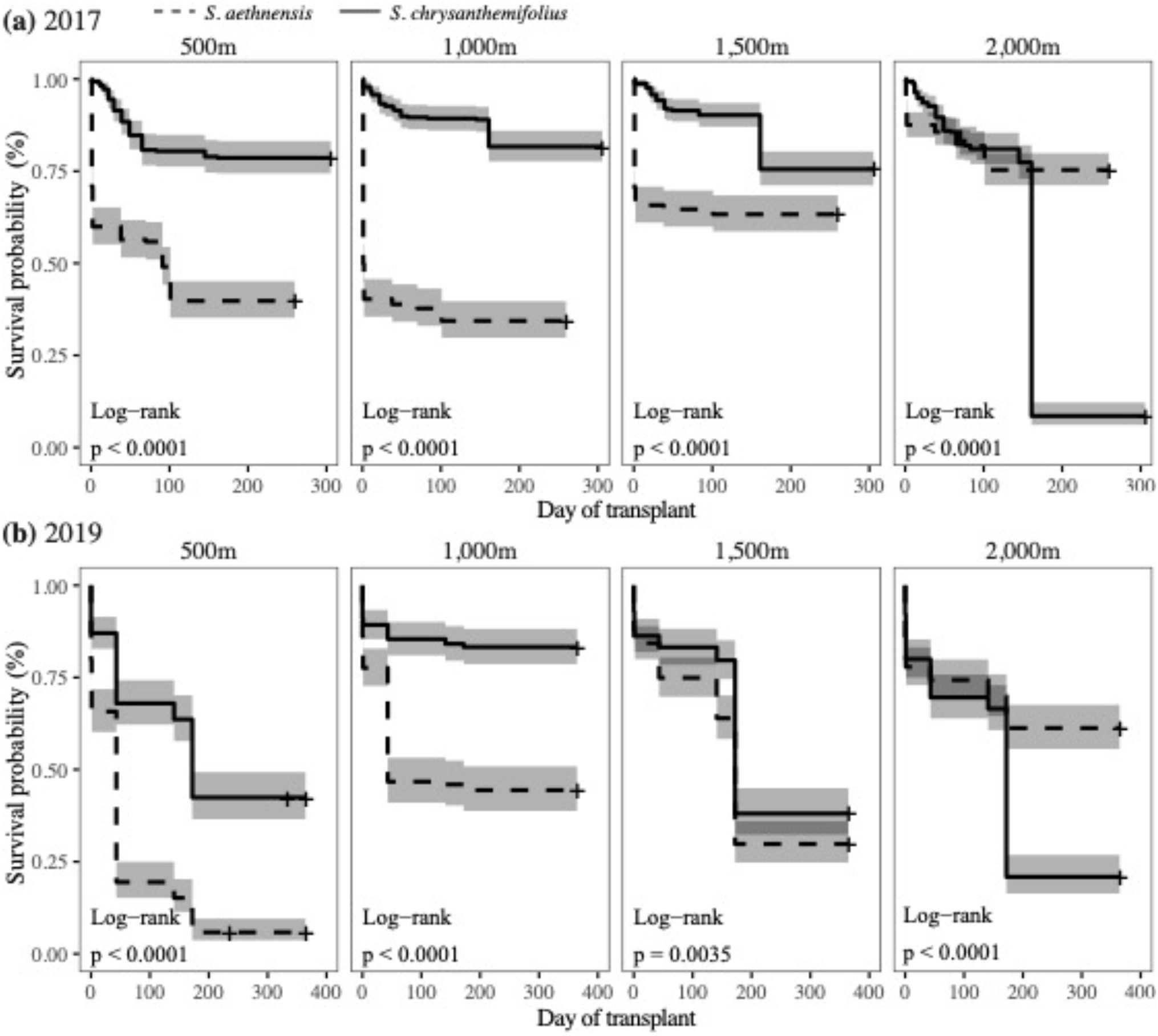
Differences in survival between species across all transplant elevations, for **(a)** 2017 and **(b)** 2019. Solid lines represent *S. chrysanthemifolius*, and broken lines *S. aethnensis*. Grey shading represents the 95% confidence interval around the mean survival of each species. *Senecio chrysanthemifolius* showed the lowest survival at the highest elevation in both experiments, while *S. aethnensis* showed lower survival at all three lower elevations.

This is likely to be due to cuttings suffering under intense heat without well-developed root systems, especially given they received less supplemental water compared to 2017. However, mortality in 2019 was still c.20% greater at the novel elevation (2,000m) than at 500m, indicating better performance at the native site.

### Species differences in plasticity: Univariate trait changes across elevation

Both species showed phenotypic plasticity as a change in trait mean with elevation for most leaf morphology traits **(Fig. 5)**, with patterns of plasticity that were consistent across both experiments. However, the degree to which plastic changed the phenotype across elevation differed between species depended on the trait. Both species showed similar increases in leaf area at lower elevations (**Fig. 5a-b**; species×elevation in 2017: *χ^2^*(3)=8.94, adj. P=0.1202, and 2019: *χ^2^*(3)=42.08, adj. P<0.0001). However, *Senecio chrysanthemifolius* showed a reduction in leaf complexity at higher elevations, contrasting with no change across elevation in *S. aethnensis* (**Fig. 5c-d**; species×elevation in 2017: *χ^2^*(3)=29.09, adj. P<0.0001, and 2019: *χ^2^*(3)=83.87, adj. P<0.0001). By contrast, *S. aethnensis* showed a reduction in leaf indentation at higher elevations, while *S. chrysanthemifolius* showed no significant change in leaf indentation across elevation in 2017, but a significant increase in leaf indentation at 2,000m in 2019 (**Fig. 5e-f**; species×elevation in 2017: *χ^2^*(3)=29.07, adj. P<0.0001, and 2019: *χ^2^*(3)=66.68, adj. P<0.0001). Both species also showed a similar increase in SLA (Specific Leaf Area) at lower elevations, but *S. aethnensis* showed a much greater increase at 500m and 1,000m (**Fig. 5g-h**; species×elevation in 2017: *χ^2^*(3)=21.95, adj. P=0.00027, and 2019: *χ^2^*(3)=37.71, adj. P<0.0001), suggesting that *S. aethnensis* produced much thinner leaves than *S. chrysanthemifolius* at lower elevations (and at all elevations in 2019).

**Fig. 5.**
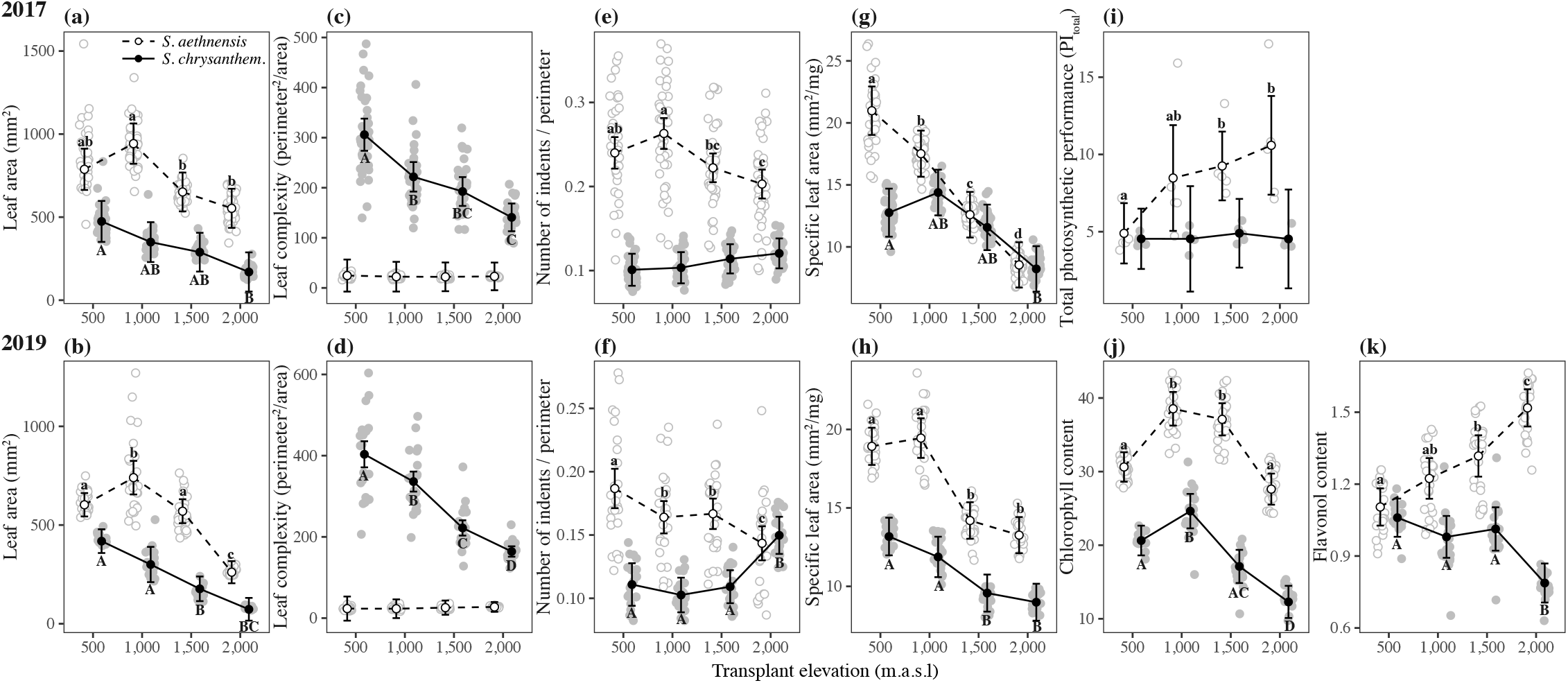
Changes in univariate leaf traits across elevation for both species measured in 2017 (top row) and 2019 (bottom row). Filled circles, solid lines and upper case letters represent *S. chrysanthemifolius*, while unfilled circles, dashed lines and lowercase letters represent *S. aethnensis*. Credible intervals represent 95% confidence intervals for the estimate of the mean of each species at each elevation. Letters denote significant differences between transplant sites calculated using pairwise tests conducted within each species and adjusted for multiple comparisons (full statistical summaries are located in **Table S3**). Grey points represent the mean of all cuttings for each genotype. Traits include: **(a-b)** leaf area, **(c-d)** leaf complexity, **(e-f)** number of indents, **(g-h)** specific leaf area, **(i)** total photosynthetic performance, **(j)** chlorophyll content and **(k)** flavonol content. Results are consistent across the 2017 and 2019 experiments, and show that the two species display distinct plastic responses across elevation for leaf complexity, number of indents, specific leaf area, photosynthetic performance and flavonol content.

In 2017, we measured chlorophyll fluorescence to calculate the total performance index (PI_total_), which estimates the capacity of the photosynthetic machinery. Although *S. chrysanthemifolius* showed no change in PI_total_ across elevation, *S. aethnensis* showed a steady decline, suggesting reduced photosynthetic activity of *S. aethnensis* at lower elevations (**Fig. 5i**; species×elevation *χ^2^*(3)=24.59, P<0.0001). In 2019, we measured the leaf concentration of chlorophyll and flavonols. Compared to *S. aethnensis, S. chrysanthemifolius* showed higher concentrations of both pigments. From high to low elevation, both species showed increases in chlorophyll content, but a reduction at 500m (**Fig. 5j**; species×elevation: *χ^2^*(3)=64.23, adj. P<0.0001). The two species show contrasting patterns of plasticity in flavonol content across elevation (**Fig. 5k**; species×elevation: *χ^2^*(3)=98.57, adj. P<0.0001): although both species show similar values at 500m, flavonol content in *S. chrysanthemifolius* decreases at higher elevations, while in *S. aethnensis* it increases at higher elevations.

### Species differences in plasticity: Change in multivariate phenotype across elevation

To estimate multivariate plasticity, we quantified variation in leaf morphology between species and across elevation by analysing all leaf traits using a multivariate analysis of variance (MANOVA). For both experiments, we found significant species×elevation interactions (**Fig. 6;** 2017: Wilks’ λ=0.375, *F*_3,298_=29.19, P<0.0001; 2019: Wilks’ λ=1.330, *F*_3,176_=22.97, P<0.0001). In both experiments, the first axis of **D**(***d***_max_) described >75% of the difference in multivariate phenotype, which primarily separated the two species, generated by a negative correlation between leaf area and complexity (larger simple leaves versus smaller, more complex leaves). The second axis (***d***_2_) described 15-19% of the difference in multivariate phenotype, which captured differences across elevation as reductions in most traits at higher elevations. Outside its native range (i.e. at 500m), plasticity moved the multivariate phenotype of *S. aethnensis* away from, rather than towards, the native phenotype of *S. chrysanthemifolius.* By contrast, *S. chrysanthemifolius* genotypes outside their native range (i.e. at 2000m) shifted their phenotypes towards that expressed by *S. aethnensis* at high elevations (**Fig. 6**). The vectors representing elevational plasticity 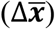 differed between species by 54° and 56° respectively for the 2017 and 2019 experiments (grey lines in **Fig. 6**), which suggests the two species showed plastic responses to elevation that differ consistently in direction across years.

**Fig. 6.**
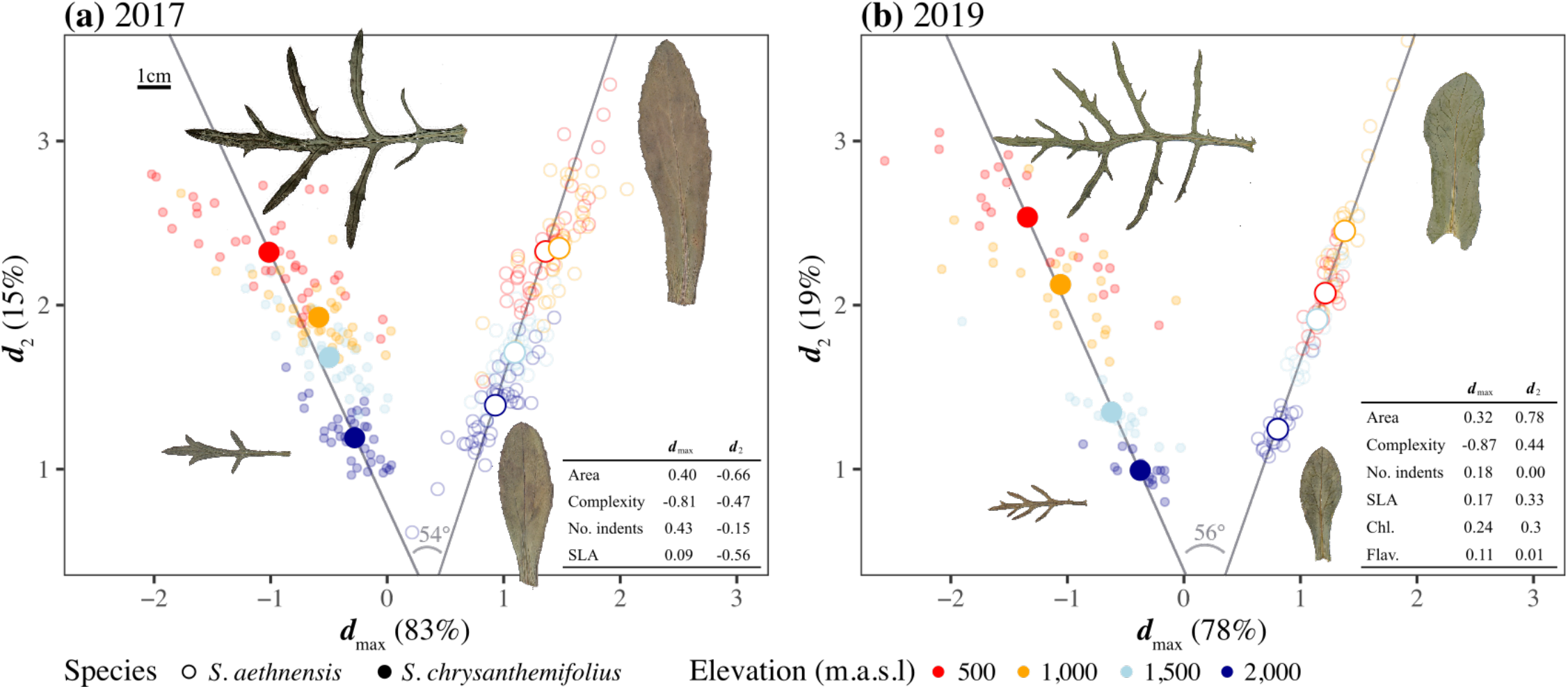
Multivariate analysis (MANOVA used to calculate the D-matrix) of leaf morphology across elevation for both species measured in **(a)** 2017 and **(b)** 2019. For both experiments, ***d***_max_ represents a trade-off between leaf size and complexity, and separates both species. By contrast, ***d***_2_ represents similar elevational changes in most traits. Small filled circles represent genotypes of *S. chrysanthemifolius*, and unfilled circles *S. aethnensis*. Large circles represent the multivariate mean at each elevation. Table inset shows the trait loadings for both multivariate axes. Inset leaf images represent a genotype of each species with a value close to the multivariate mean at the elevational extremes (500m and 2,000m). Gray lines represent the vector of plastic responses 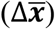 across elevation for each species. The angle between plasticity vectors show that the two species differ in the direction of plasticity by c.55°, suggesting distinct patterns of plasticity between species.

### Differential gene expression between species

Comparisons of gene expression between species for the 2019 transplant revealed that many genes were differentially expressed between species at each transplant site (**Fig. 7a**). In total, the largest number of differentially expressed genes between the two species was at 1,500m (332 genes) with the fewest at 500m (279 genes). Despite observing the largest number of differentially expressed genes at 1,500m, functional enrichment revealed that the two species show the fewest significantly enriched GO terms (9 terms) at 1,500m while the largest number of significantly enriched GO terms was at 2,000m (16 terms). Furthermore, 140 genes were differentially expressed between species at all transplant sites, which represents 42-50% of the total number of differentially expressed genes at each site (and 29% of all genes differentially expressed at all elevations). Therefore, almost half the genes that were differentially expressed between species remained consistent across elevation. Despite this, no GO terms were significantly enriched between species at all transplant sites (**Fig. 7b**).These results suggest that although some genes are consistently differentially expressed between the two species, they are not related to known functional categories. This means that the functional categories of genes showing large differences in gene expression between species at each elevation are not shared across elevations, indicating that the two species differ in gene expression profiles in response to elevation.

**Fig. 7.**
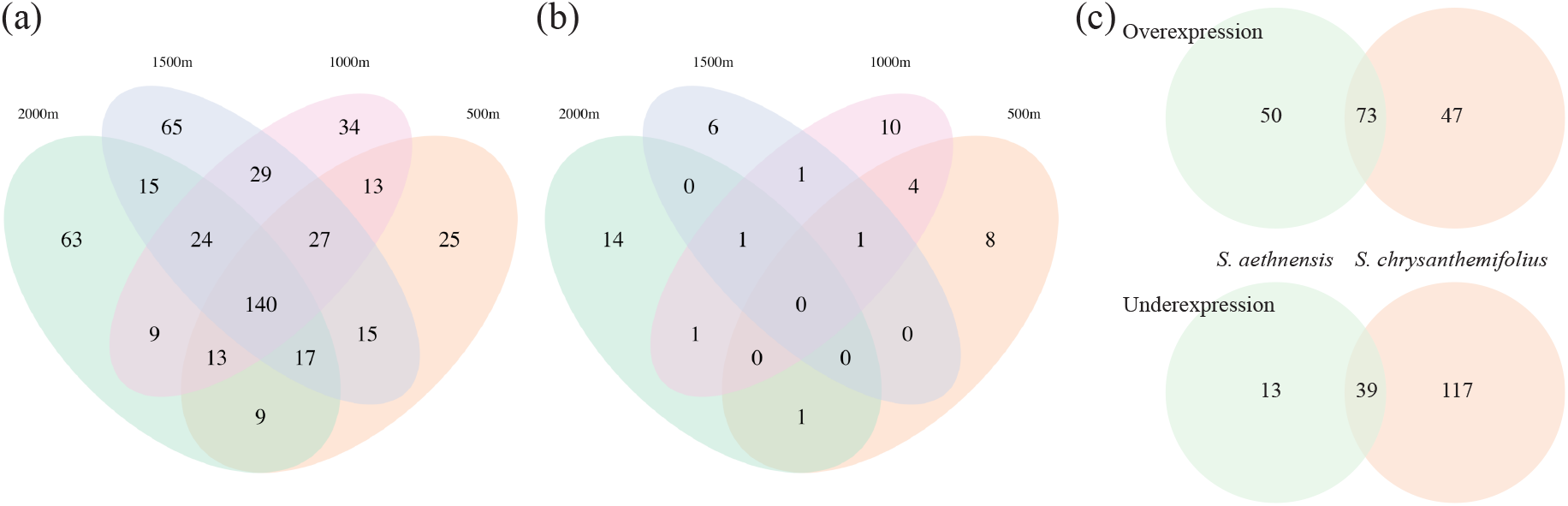
Contrasting patterns of gene expression plasticity between species. (**a**) Total numbers of differentially expressed genes (−2 < lfc > 2) between species at each transplant site. Many genes that were differentially expressed between the species were unique to each elevation. **(b)** Total numbers of significantly enriched (Fisher’s exact test, p < 0.05) GO terms between species at each transplant site. Only found significant functional categories were found for differentially expressed genes at each elevation, which were rarely shared across elevations. (**c**) Overlapping overexpressed and underexpressed genes between the home and furthest transplant site in each species. Many genes that were differentially expressed across elevation were unique to each species.

### Differential gene expression across elevation within species

The two methods of estimating differential expression (*DESeq2* vs *limma/voom*) showed similar patterns of gene expression variation, although *limma/voom* detected fewer strongly differentially expressed genes in each species (**Fig. S1**). For both species, more genes were differentially expressed as genotypes were moved further from their native elevations (**Fig. S1)**. We identified c.350 loci (out of c. 19,000) per species that showed differential expression between the extreme elevations. Only a small proportion of genes that were differentially expressed between the native site and the most novel environment (i.e. the elevational extremes) were common to both species, with 43% and 23 % respectively of overexpressed and underexpressed genes showing similar patterns across elevation in both species (**Fig. 7c**). However, the majority of differentially expressed genes showed a species-specific response to elevation providing evidence for distinct patterns of plasticity in gene expression for the two species (**Fig. 7c**). This meant that most genes showed responses to elevation that were distinct to each species: strong overexpression or underexpression in one species contrasted with relatively unchanged expression across elevation in the other species (**Fig. 8**). Functional enrichment analyses of differentially expressed genes between the elevational extremes revealed 9 significant GO terms in *S. chrysanthemifolius* and 20 in *S. aethnensis*. Only 2 functional categories of these genes were shared between species, again suggesting that many genes responded in a species-specific manner (**Tables S4-S5**).

**Fig. 8.**
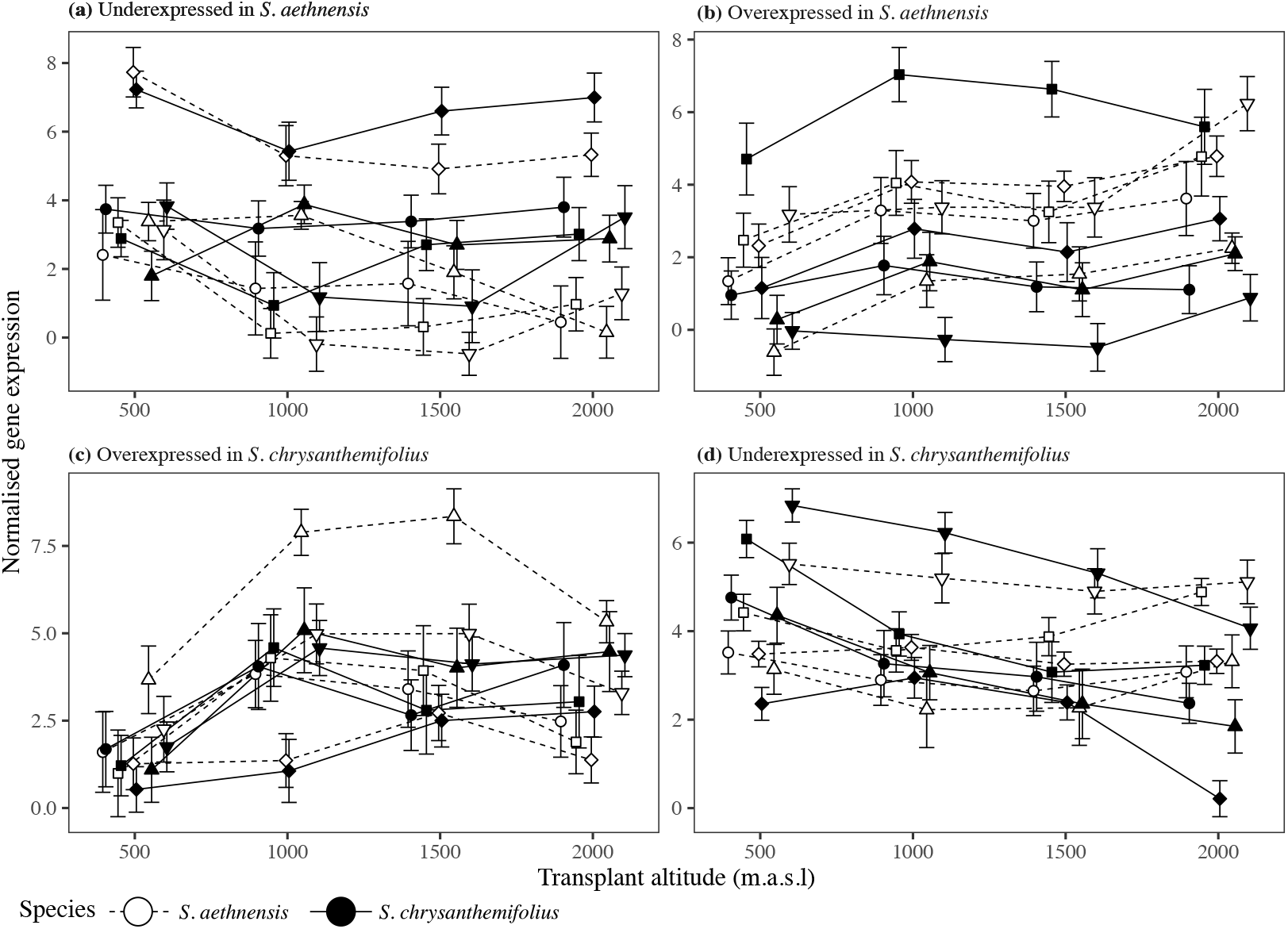
Strong overexpression or underexpression in one species was typically reflected by little to no change in gene expression in the same gene for the other species. Normalised expression profiles across all transplant sites for *S. chrysanthemifolius* (solid lines and circles) compared to *S. aethnensis* (dashed lines and unfilled circles). This includes the genes (represented by different shapes) that most strongly underexpressed in *S. aethnensis* but not in *S. chrysanthemifolius* **(a)**, overexpressed in *S. aethnensis* but not in *S. chrysanthemifolius* **(b)**, and five genes strongly underexpressed in *S. chrysanthemifolius* but not in *S. aethnensis* **(c)** and overexpressed in *S. chrysanthemifolius* but not in *S. aethnensis* **(d)**.

Network reconstruction generated 13 network modules ranging in size from 185 to 3345 genes (**Table S6**). In 6 modules the module eigengene correlated with elevation in both species and 2 modules (**Table S6**: GN3 and GN8) showed a correlation with elevation unique to *S. aethnensis*. No gene modules that correlated with elevation were specific to *S. chrysanthemifolius.* The two modules that changed in *S. aethnensis* but not *S. chrysanthemifolius* were associated with functions related to pathogen detection and responses to light intensity and UV.

## Discussion

Given the two *Senecio* species on Mt. Etna are the result of recent ecological divergence (Chapman et al. 2013), we predicted that: (1) the two species would show differences in physiology when grown in common garden conditions; (2) they would perform better at their native elevations than beyond their existing range, and better than the other species within their native habitat; and (3) given their adaptation to contrasting environments, they would display different patterns of plasticity at the phenotypic and gene expression levels.

Consistent with our first prediction, both species showed distinct behaviour in the common garden experiment (**Fig. 2**), indicating that adaptation to contrasting habitats has generated substantial differences in physiology between the two species. Supporting our second prediction, both species performed better at their native site than the other species. Each species also tended to perform best at their native site compared to other elevations, with strong reductions in survival observed at the elevation furthest from their native site (i.e. at the native site of the other species; **Fig. 4**). This provides strong evidence that these two species are adapted to their natural habitats, which is consistent with another field experiment that demonstrated adaptive divergence between these two species when transplanted as seedlings rather than as cuttings (Walter et al. 2021). Consistent with our third prediction, we found different patterns of plasticity in phenotype (**Figs. 5-6)** and gene expression profiles (**Figs. 7-8**) between the two species.

### Species differences in gene expression plasticity

The native conditions experienced by any given species can determine how gene expression will change in response to environmental variation (Akman et al. 2016). We observed both differential expression between species as well as between-species differences in the plasticity of gene expression along the elevational gradient. Although differential expression between species was observed at each transplant site, we found that only 29% (140 genes) of genes that were differentially expressed between species were common to all transplant sites. This suggests that each species has recently evolved distinct patterns of gene expression associated with their native habitats.

Within a species, plasticity in gene expression in this experiment is represented by genes that are differentially expressed between their native elevation and other elevations. We found that only 33% of genes that were differentially expressed across elevational extremes were shared by the two species. The differences in plasticity between species are evident in the expression profiles of individual genes, where genes that showed strong changes in one species were weaker or absent in the other species **(Fig 8)**. This meant that highly divergent patterns of gene expression between the species were rare and expression profiles tended to reflect differences in the magnitude of the response between species, rather than responses in opposite directions (**Fig 8**). We also found unique networks associated with the response of *S. aethnensis* to elevation, suggesting that differential expression in specific gene networks underlies differences in plasticity observed between the two species, particularly in response to biotic stressors and changes in light conditions associated with higher elevation. Together, these results confirm that these species show distinct patterns of gene expression responses to elevation, most often arising from species-specific patterns of expression at similar loci, but with some loci and gene networks displaying contrasting patterns of expression between the species.

### Adaptive divergence creates differences in plasticity

Our results suggest that adaptive divergence between these species is not only associated with divergence in mean phenotype and physiology under common garden conditions, but also divergence in patterns of phenotypic plasticity and in the genetic basis of plasticity. It remains possible that genetic drift during the formation of these two species could cause the species-specific differences in plasticity that we observed. However, given substantial divergence in leaf form and physiology under common garden conditions, the differences in plasticity of these same traits among species, and the higher fitness of each species at their native versus novel habitats, it seems far more likely that adaptive divergence between *S. aethnensis* and *S. chrysanthemifolius* is responsible for their distinct plastic responses (Taylor and Aarssen 1988; Emery et al. 1994; Donohue et al. 2001; Ho and Zhang 2018).

Other studies have shown that species from different genera can evolve distinct forms of plasticity (Schlichting and Levin 1986). One study that compared closely related species of *Phlox* wildflowers found inconsistent differences that could not be explained by ecology or relatedness (Schlichting and Levin 1984). Here, by comparing plasticity for two closely related species in field experiments that include their natural habitats, we show that adaptive divergence is associated with the evolution of different forms of plasticity. We also show that the magnitude and direction of plasticity in response to elevation is trait dependent, in the sense that the two species showed differences in the magnitude of plasticity across elevation in some traits (e.g. leaf complexity and specific leaf area), and a contrasting pattern of plasticity in other traits (e.g. leaf indents and flavonol content). This suggests that adaptive divergence can change both the magnitude and direction of plasticity, which is likely to depend on how selection varies its action on different traits (Via 1993).

### An ecological explanation for differences in plasticity

As a consequence of adapting to the environmental variation specific to their native elevations, these species could show different patterns of plasticity because they only maintain plasticity in traits important for tracking variation within their native environment. For example, as a perennial that grows from rootstock after winter each year, it is likely that the new leaves *S. aethnensis* produces each year possess different characteristics that are optimal for the specific environmental conditions of that year, allowing *S. aethnensis* to track environmental variation across years. In this case, plasticity in leaf traits would be critical to cope with environmental variation across years at high elevations, and so would also show plasticity when transplanted to lower elevations, even if such plastic changes were maladaptive. This would mean that how selection acts on a trait in different environments will likely determine the levels of plasticity that evolves in populations adapting to each environment.

As divergent selection drives trait divergence between the two species, it will likely also change the level of plasticity in the traits that are diverging. Species will therefore show reduced plasticity in the genes (i.e. the genes that show differential expression across elevations) and in the traits that need to be continually expressed in their native habitat. This would occur if the plastic responses required to initially colonise a high or low elevation habitat become genetically fixed (via genetic assimilation) because consistently strong stabilising selection for a single phenotype is favoured (Waddington 1953). In this scenario, plasticity would only be retained in traits that maintain fitness in response to the environmental variation experienced within their native habitat. For example, *S. chrysanthemifolius* shows both greater leaf complexity than *S. aethnensis*, as well as greater plasticity in this trait (**Fig. 5b & 5g**), while *S. aethnensis* shows a greater number of indents, as well as greater plasticity in this trait (**Fig. 5c & 5h**). At high elevations *Senecio aethnensis* is likely to experience consistently strong selection for less complex leaves, leading to a loss of plasticity in this trait, perhaps via genetic assimilation. By contrast, *S. chrysanthemifolius* is likely to experience selection for more complex leaves, and the greater variety of habitats that this species occupies could maintain plasticity in this trait. Future work should identify how selection affects the evolution of plasticity in different traits and environmental regimes (Schmitt et al. 1999; Pratt and Mooney 2013; McLean et al. 2014), and could reveal how the fixation of alleles during adaptation affects trait plasticity (Schlichting and Levin 1986; Forsman 2015).

### Predicting species responses to novel environments

Understanding how plasticity evolves is important for understanding how species can respond to novel environmental variation (Bradshaw 1965; Baythavong and Stanton 2010). For example, plants experiencing strong stabilizing selection for a particular phenotype will become more specialised and evolve reduced plasticity (Emery et al. 1994; Alpert and Simms 2002; Baythavong 2011). In other studies, high-elevation populations showed reduced plasticity in flowering time (Schmid et al. 2017) and morphology (Emery et al. 1994) compared to low elevation populations. Similarly, we found that the high elevation species, *S. aethnensis*, showed a steep decline in survival (**Fig. 4**), which was associated with stronger reductions in leaf investment (**Fig. 5d** & **6i**), lower photosynthetic activity (**Fig. 5e**) and reduced capacity for plasticity in leaf traits to produce phenotypes more similar to the native phenotype of *S. chrysanthemifolius* at lower elevations (**Fig. 6**). Therefore, the contrasting patterns of plasticity observed for the two species may be caused by *S. aethnensis* exhibiting maladaptive plasticity at lower elevations (1,500m and below) as a consequence of adapting to the high elevation habitat.

Limits to adaptive plasticity could threaten the persistence of populations exposed to novel environments unless genotypic variation in plasticity (genotype×environment interactions) can allow rapid adaptation (Lande 2009; Chevin et al. 2010). Understanding whether genetic variation in plasticity will allow natural populations to adapt to ongoing environmental change requires quantifying plasticity and fitness across environmental variation for genotypes from natural populations.

