## Supplementary material for "Adaptive divergence generates distinct plastic responses in two closely related *Senecio* species"

**Contents**

**Table S1** Sampling locations

**Table S2** Soil analysis

**Table S3** Statistical output of the pairwise comparisons for all morphological traits

**Fig. S1** Number of differentially expressed genes

**Table S4** Network analysis

**Table S5** Gene ontology for *S. aethnensis*

**Table S6** Gene ontology for *S. chrysanthemifolius*

**Methods S1** Supplemental methods

**Table S1:** Location of sampled individuals used in the transplant. Sp. = Species (*S.ae* = *S. aethnensis; S.ch = S. chrysanthemifolius*), Elev. = Elevation, ind. = individuals. Clones/site represent the number of clones transplanted at each field transplant site. For the 2017 experiment, individuals varied in the number of cuttings that successfully produced roots in the glasshouse. Consequently, we transplanted individuals with 6, 9 or 15 clones at each transplant site, depending on the availability of cuttings. Columns denoting the number of clones transplanted per site refer to how many genotypes in each category, and from which site in the natural populations those genotypes were sampled in 2017. The final column denotes the number of maternal genotypes (from each site) that seeds were collected and germinated to grow the genotypes for the 2019 transplant.

|  |  |  |  |  | **2017 transplant experiment** | | | |  | **2019** |
| --- | --- | --- | --- | --- | --- | --- | --- | --- | --- | --- |
| **Sp.** | **Site** | **Elev.** | **Latitude** | **Longitude** | **6 clones/**  **site** | **9 clones/**  **site** | **15 clones/**  **site** | **Total ind.** |  | **Maternal genotypes** |
| *S. ae* | Etna South | 2,600m | 37°43'13.28"N | 15° 0'3.54"E | 2 | 1 | 0 | 3 |  | 4 |
|  |  | 2,500m | 37°43'3.80"N | 14°59'59.20"E | 3 | 4 | 2 | 9 |  | 3 |
|  |  | 2,400m | 37°42'46.50"N | 14°59'41.30"E | 3 | 2 | 1 | 6 |  | 5 |
|  |  | 2,200m | 37°42'24.82"N | 14°59'42.69"E | 2 | 2 | 0 | 4 |  | 1 |
|  | Etna North | 2,600m | 37°46'39.90"N | 15° 0'23.00"E | 0 | 5 | 1 | 6 |  | 3 |
|  |  | 2,500m | 37°46'53.70"N | 15° 0'28.80"E | 1 | 2 | 2 | 5 |  | 3 |
|  |  | 2,400m | 37°47'7.46"N | 15° 0'35.05"E | 2 | 2 | 2 | 6 |  | 3 |
|  |  | 2,200m | 37°47'32.82"N | 15° 1'14.53"E | 1 | 1 | 1 | 3 |  | 3 |
|  | **Totals** |  |  |  | **14** | **19** | **9** | **42** |  | **25** |
| *S. ch* | Bonnano | 790m | 37°38'24.92"N | 15° 2'50.80"E | 4 | 2 | 0 | 6 |  | 3 |
|  | Cacciola | 680m | 37°37'31.32"N | 15° 3'26.71"E | 4 | 3 | 3 | 10 |  | 5 |
|  | Poggofelice | 590m | 37°39'44.31"N | 15° 5'48.55"E | 3 | 3 | 1 | 7 |  | 5 |
|  | Spina | 730m | 37°39'19.27"N | 15° 4'30.92"E | 4 | 3 | 3 | 10 |  | 5 |
|  | Trecastagni | 570m | 37°36'46.67"N | 15° 4'29.64"E | 3 | 0 | 1 | 4 |  | 3 |
|  | **Totals** |  |  |  | **18** | **11** | **8** | **37** |  | **21** |

**Table S2** Analysis of the soil samples

| Elevation | Replicate | Total N | C:N | Organic matter | Available P | Available K | Total K | pH | Conductibility | Soluble Chloride | Soluble nitrate | Soluble sulphites |
| --- | --- | --- | --- | --- | --- | --- | --- | --- | --- | --- | --- | --- |
|  |  | g/kg |  | g/kg | mg/kg | mg/kg | mg/kg |  | dS/m | meq/L | meq/L | meq/L |
| 500m | A | 1.4 | 9.5 | 23 | 57 | 122 | 9850 | 7.7 | 0.31 | 0.8 | 0 | 1.2 |
| 500m | B | 1.3 | 8.9 | 20 | 57 | 165 | 9347 | 7.5 | 0.25 | 0.5 | 0.7 | 0.3 |
| 500m | C | 1.1 | 7.1 | 13 | 39 | 158 | 9499 | 7.6 | 0.21 | 0.8 | 0 | 0.6 |
| 1000m | A | 0.9 | 8.6 | 13 | 14 | 57 | 10160 | 7.1 | 0.45 | 0.3 | 3.3 | 0.2 |
| 1000m | B | 1.2 | 8.2 | 17 | 14 | 70 | 10240 | 7 | 0.54 | 0.4 | 4.1 | 0.3 |
| 1000m | C | 1 | 7.2 | 12 | 11 | 51 | 9631 | 7.2 | 0.54 | 0.5 | 3.7 | 0.3 |
| 1500m | A | 0.9 | 5.6 | 9 | 16 | 72 | 9333 | 7.2 | 0.16 | 0.5 | 0.3 | 0.3 |
| 1500m | B | 0.9 | 4.7 | 7 | 16 | 92 | 9520 | 7.2 | 0.15 | 0.5 | 0.6 | 0.2 |
| 1500m | C | 1 | 5.6 | 10 | 21 | 68 | 9265 | 7.2 | 0.26 | 0.2 | 1.6 | 0.2 |
| 2000m | A | 0.8 | 6.1 | 8 | 69 | 83 | 10680 | 8 | 0.38 | 0.8 | 1.1 | 0.9 |
| 2000m | B | 0.9 | 2.3 | 4 | 14 | 48 | 11460 | 7.2 | 0.14 | 0.3 | 0.3 | 0.2 |
| 2000m | C | 0.7 | 3 | 4 | 18 | 83 | 11240 | 7.3 | 0.12 | 0.5 | 0 | 0.4 |
| Elevation | Replicate | Soluble sodium | Soluble calcium | Soluble magnesium | Soluble potassium | Cation exchange capacity | Exchangeable calcium | Exchangeable magnesium | Exchangeable sodium | Exchangeable potassium | K:Mg |  |
|  |  | meq/L | meq/L | meq/L | meq/L | meq/100g | meq/100g | meq/100g | meq/100g | meq/100g |  |  |
| 500m | A | 0.4 | 0.8 | 0.6 | 0.2 | 5.6 | 3.9 | 1 | 0.5 | 0.3 | 0.26 |  |
| 500m | B | 0.3 | 0.4 | 0.7 | 0.2 | 5.7 | 4.2 | 0.7 | 0.4 | 0.4 | 0.5 |  |
| 500m | C | 0.3 | 0.4 | 0.5 | 0.2 | 5 | 3.5 | 0.8 | 0.4 | 0.3 | 0.44 |  |
| 1000m | A | 0.6 | 1.6 | 1.1 | 0.3 | 3.2 | 2.4 | 0.4 | 0.3 | 0.1 | 0.3 |  |
| 1000m | B | 0.9 | 2 | 1.3 | 0.3 | 4.4 | 2.9 | 0.5 | 0.8 | 0.1 | 0.28 |  |
| 1000m | C | 0.9 | 2.1 | 1 | 0.3 | 2.8 | 2.1 | 0.3 | 0.3 | 0.1 | 0.34 |  |
| 1500m | A | 0.3 | 0.3 | 0.4 | 0.4 | 3.4 | 2.6 | 0.4 | 0.2 | 0.2 | 0.42 |  |
| 1500m | B | 0.3 | 0.2 | 0.4 | 0.2 | 2.6 | 1.8 | 0.4 | 0.2 | 0.2 | 0.54 |  |
| 1500m | C | 0.5 | 0.5 | 0.6 | 0.2 | 2.7 | 1.9 | 0.3 | 0.3 | 0.1 | 0.43 |  |
| 2000m | A | 0.5 | 1.5 | 0.8 | 0.2 | 5.3 | 4.3 | 0.5 | 0.3 | 0.2 | 0.34 |  |
| 2000m | B | 0.3 | 0.2 | 0.2 | 0 | 2.7 | 1 | 0.2 | 1.4 | 0.1 | 0.43 |  |
| 2000m | C | 0.4 | 0.2 | 0.2 | 0.1 | 2.3 | 1.5 | 0.4 | 0.2 | 0.2 | 0.45 |  |

**Table S3** Full statistical summaries of each trait for the post-hoc pairwise comparison of transplant sites for each species (*S.ae* = *S. aethnensis; S.ch = S. chrysanthemifolius*).

| **Trait** | **contrast** | **Species** | **Estimate** | **SE** | **df** | **z-ratio** | **Adj. P-value** |
| --- | --- | --- | --- | --- | --- | --- | --- |
| **Fig. 4a** 2017 survival | 500m - 1000m | *S.ae* | -0.373 | 0.328 |  | -1.136 | 0.667 |
|  | 500m - 1500m | *S.ae* | 0.642 | 0.333 |  | 1.929 | 0.216 |
|  | 500m - 2000m | *S.ae* | 1.312 | 0.338 |  | 3.878 | 0.001 |
|  | 1000m - 1500m | *S.ae* | 1.015 | 0.332 |  | 3.054 | 0.012 |
|  | 1000m - 2000m | *S.ae* | 1.685 | 0.338 |  | 4.985 | 0.000 |
|  | 1500m - 2000m | *S.ae* | 0.670 | 0.342 |  | 1.958 | 0.204 |
|  | 500m - 1000m | *S.ch* | 0.218 | 0.364 |  | 0.598 | 0.933 |
|  | 500m - 1500m | *S.ch* | 0.031 | 0.361 |  | 0.086 | 1.000 |
|  | 500m - 2000m | *S.ch* | -1.665 | 0.342 |  | -4.862 | 0.000 |
|  | 1000m - 1500m | *S.ch* | -0.187 | 0.365 |  | -0.512 | 0.956 |
|  | 1000m - 2000m | *S.ch* | -1.882 | 0.347 |  | -5.426 | 0.000 |
|  | 1500m - 2000m | *S.ch* | -1.696 | 0.344 |  | -4.933 | 0.000 |
| **Fig. 4b**  2019 survival | 500m - 1000m | *S.ae* | 1.293 | 0.429 |  | 3.010 | 0.014 |
|  | 500m - 1500m | *S.ae* | 1.146 | 0.427 |  | 2.682 | 0.037 |
|  | 500m - 2000m | *S.ae* | 1.887 | 0.433 |  | 4.356 | 0.000 |
|  | 1000m - 1500m | *S.ae* | -0.146 | 0.431 |  | -0.340 | 0.987 |
|  | 1000m - 2000m | *S.ae* | 0.594 | 0.436 |  | 1.363 | 0.522 |
|  | 1500m - 2000m | *S.ae* | 0.741 | 0.434 |  | 1.707 | 0.320 |
|  | 500m - 1000m | *S.ch* | 1.583 | 0.454 |  | 3.484 | 0.003 |
|  | 500m - 1500m | *S.ch* | 0.027 | 0.433 |  | 0.063 | 1.000 |
|  | 500m - 2000m | *S.ch* | -0.431 | 0.431 |  | -1.001 | 0.749 |
|  | 1000m - 1500m | *S.ch* | -1.555 | 0.454 |  | -3.427 | 0.003 |
|  | 1000m - 2000m | *S.ch* | -2.014 | 0.452 |  | -4.455 | 0.000 |
|  | 1500m - 2000m | *S.ch* | -0.459 | 0.430 |  | -1.066 | 0.710 |
| **Fig. 5a**  2017 leaf area | 500m - 1000m | *S.ae* | -154.800 | 79.384 | 17.587 | -1.950 | 0.244 |
|  | 500m - 1500m | *S.ae* | 136.388 | 79.367 | 17.583 | 1.718 | 0.344 |
|  | 500m - 2000m | *S.ae* | 234.839 | 80.012 | 18.152 | 2.935 | 0.040 |
|  | 1000m - 1500m | *S.ae* | 291.188 | 78.645 | 16.970 | 3.703 | 0.009 |
|  | 1000m - 2000m | *S.ae* | 389.638 | 79.238 | 17.484 | 4.917 | 0.001 |
|  | 1500m - 2000m | *S.ae* | 98.451 | 77.519 | 16.037 | 1.270 | 0.594 |
|  | 500m - 1000m | *S.ch* | 124.511 | 77.906 | 16.367 | 1.598 | 0.407 |
|  | 500m - 1500m | *S.ch* | 185.272 | 78.624 | 16.962 | 2.356 | 0.124 |
|  | 500m - 2000m | *S.ch* | 304.482 | 79.540 | 17.742 | 3.828 | 0.006 |
|  | 1000m - 1500m | *S.ch* | 60.762 | 77.739 | 16.225 | 0.782 | 0.862 |
|  | 1000m - 2000m | *S.ch* | 179.971 | 78.606 | 16.940 | 2.290 | 0.140 |
|  | 1500m - 2000m | *S.ch* | 119.209 | 77.481 | 15.988 | 1.539 | 0.439 |

**Table S3** (cont’d)

| **Trait** | **contrast** | **Species** | **Estimate** | **SE** | **df** | **t ratio** | **P-value** |
| --- | --- | --- | --- | --- | --- | --- | --- |
| **Fig. 5b**  2019 leaf area | 500m - 1000m | *S.ae* | -137.354 | 45.159 | 28.899 | -3.042 | 0.024 |
|  | 500m - 1500m | *S.ae* | 32.617 | 37.760 | 16.201 | 0.864 | 0.823 |
|  | 500m - 2000m | *S.ae* | 341.633 | 38.086 | 16.634 | 8.970 | 0.000 |
|  | 1000m - 1500m | *S.ae* | 169.972 | 41.696 | 22.846 | 4.076 | 0.002 |
|  | 1000m - 2000m | *S.ae* | 478.988 | 47.726 | 33.067 | 10.036 | 0.000 |
|  | 1500m - 2000m | *S.ae* | 309.016 | 38.280 | 17.013 | 8.073 | 0.000 |
|  | 500m - 1000m | *S.ch* | 118.320 | 46.307 | 29.982 | 2.555 | 0.072 |
|  | 500m - 1500m | *S.ch* | 241.556 | 38.240 | 16.709 | 6.317 | 0.000 |
|  | 500m - 2000m | *S.ch* | 345.330 | 38.984 | 17.800 | 8.858 | 0.000 |
|  | 1000m - 1500m | *S.ch* | 123.236 | 42.623 | 23.985 | 2.891 | 0.038 |
|  | 1000m - 2000m | *S.ch* | 227.011 | 49.841 | 35.681 | 4.555 | 0.000 |
|  | 1500m - 2000m | *S.ch* | 103.774 | 39.563 | 18.794 | 2.623 | 0.073 |
| **Fig. 5c**  2017 leaf complexity | 500m - 1000m | *S.ae* | 2.241 | 19.131 | 19.651 | 0.117 | 0.999 |
|  | 500m - 1500m | *S.ae* | 2.546 | 18.709 | 18.011 | 0.136 | 0.999 |
|  | 500m - 2000m | *S.ae* | 1.655 | 19.277 | 20.229 | 0.086 | 1.000 |
|  | 1000m - 1500m | *S.ae* | 0.305 | 18.382 | 16.813 | 0.017 | 1.000 |
|  | 1000m - 2000m | *S.ae* | -0.586 | 18.669 | 17.869 | -0.031 | 1.000 |
|  | 1500m - 2000m | *S.ae* | -0.891 | 18.285 | 16.469 | -0.049 | 1.000 |
|  | 500m - 1000m | *S.ch* | 84.122 | 18.816 | 18.444 | 4.471 | 0.001 |
|  | 500m - 1500m | *S.ch* | 113.390 | 18.548 | 17.428 | 6.113 | 0.000 |
|  | 500m - 2000m | *S.ch* | 165.014 | 19.220 | 19.993 | 8.586 | 0.000 |
|  | 1000m - 1500m | *S.ch* | 29.268 | 18.166 | 16.058 | 1.611 | 0.400 |
|  | 1000m - 2000m | *S.ch* | 80.892 | 18.521 | 17.315 | 4.368 | 0.002 |
|  | 1500m - 2000m | *S.ch* | 51.624 | 18.275 | 16.420 | 2.825 | 0.053 |
| **Fig. 5d**  2019 Leaf complexity | 500m - 1000m | *S.ae* | 0.309 | 9.281 | 37.208 | 0.033 | 1.000 |
|  | 500m - 1500m | *S.ae* | -2.371 | 10.968 | 45.335 | -0.216 | 0.996 |
|  | 500m - 2000m | *S.ae* | -4.054 | 13.195 | 49.119 | -0.307 | 0.990 |
|  | 1000m - 1500m | *S.ae* | -2.680 | 8.332 | 29.800 | -0.322 | 0.988 |
|  | 1000m - 2000m | *S.ae* | -4.363 | 10.201 | 42.017 | -0.428 | 0.973 |
|  | 1500m - 2000m | *S.ae* | -1.682 | 7.975 | 26.194 | -0.211 | 0.997 |
|  | 500m - 1000m | *S.ch* | 67.208 | 9.566 | 36.271 | 7.025 | 0.000 |
|  | 500m - 1500m | *S.ch* | 181.331 | 11.607 | 44.971 | 15.623 | 0.000 |
|  | 500m - 2000m | *S.ch* | 239.350 | 14.163 | 48.651 | 16.900 | 0.000 |
|  | 1000m - 1500m | *S.ch* | 114.123 | 8.582 | 30.105 | 13.298 | 0.000 |
|  | 1000m - 2000m | *S.ch* | 172.142 | 10.799 | 42.880 | 15.940 | 0.000 |
|  | 1500m - 2000m | *S.ch* | 58.019 | 8.432 | 29.107 | 6.881 | 0.000 |

**Table S3** (cont’d)

| **Trait** | **contrast** | **Species** | **Estimate** | **SE** | **df** | **t ratio** | **P-value** |
| --- | --- | --- | --- | --- | --- | --- | --- |
| **Fig. 5e**  2017 Number of indents | 500m - 1000m | *S.ae* | -0.023 | 0.009 | 18.316 | -2.652 | 0.070 |
|  | 500m - 1500m | *S.ae* | 0.018 | 0.009 | 22.270 | 1.941 | 0.240 |
|  | 500m - 2000m | *S.ae* | 0.037 | 0.009 | 24.487 | 3.950 | 0.003 |
|  | 1000m - 1500m | *S.ae* | 0.041 | 0.009 | 22.500 | 4.455 | 0.001 |
|  | 1000m - 2000m | *S.ae* | 0.060 | 0.009 | 22.131 | 6.592 | 0.000 |
|  | 1500m - 2000m | *S.ae* | 0.019 | 0.009 | 19.314 | 2.195 | 0.160 |
|  | 500m - 1000m | *S.ch* | -0.002 | 0.008 | 15.369 | -0.295 | 0.991 |
|  | 500m - 1500m | *S.ch* | -0.013 | 0.009 | 20.556 | -1.455 | 0.481 |
|  | 500m - 2000m | *S.ch* | -0.019 | 0.009 | 23.870 | -2.089 | 0.185 |
|  | 1000m - 1500m | *S.ch* | -0.011 | 0.009 | 20.337 | -1.185 | 0.643 |
|  | 1000m - 2000m | *S.ch* | -0.017 | 0.009 | 20.944 | -1.892 | 0.262 |
|  | 1500m - 2000m | *S.ch* | -0.006 | 0.009 | 19.288 | -0.733 | 0.883 |
| **Fig. 5f**  2019 Number of indents | 500m - 1000m | *S.ae* | 0.023 | 0.005 | 38.206 | 4.449 | 0.000 |
|  | 500m - 1500m | *S.ae* | 0.020 | 0.005 | 39.350 | 3.740 | 0.003 |
|  | 500m - 2000m | *S.ae* | 0.043 | 0.006 | 40.122 | 7.732 | 0.000 |
|  | 1000m - 1500m | *S.ae* | -0.003 | 0.005 | 35.352 | -0.563 | 0.942 |
|  | 1000m - 2000m | *S.ae* | 0.021 | 0.005 | 39.174 | 3.831 | 0.002 |
|  | 1500m - 2000m | *S.ae* | 0.023 | 0.005 | 34.579 | 4.768 | 0.000 |
|  | 500m - 1000m | *S.ch* | 0.008 | 0.005 | 34.521 | 1.580 | 0.403 |
|  | 500m - 1500m | *S.ch* | 0.002 | 0.006 | 37.510 | 0.307 | 0.990 |
|  | 500m - 2000m | *S.ch* | -0.039 | 0.006 | 39.808 | -6.501 | 0.000 |
|  | 1000m - 1500m | *S.ch* | -0.007 | 0.005 | 33.069 | -1.315 | 0.560 |
|  | 1000m - 2000m | *S.ch* | -0.047 | 0.006 | 38.373 | -8.276 | 0.000 |
|  | 1500m - 2000m | *S.ch* | -0.041 | 0.005 | 35.976 | -7.727 | 0.000 |
| **Fig. 5g**  2017 SLA | 500m - 1000m | *S.ae* | 3.465 | 1.254 | 18.092 | 2.762 | 0.057 |
|  | 500m - 1500m | *S.ae* | 8.386 | 1.266 | 18.771 | 6.625 | 0.000 |
|  | 500m - 2000m | *S.ae* | 12.436 | 1.258 | 18.324 | 9.883 | 0.000 |
|  | 1000m - 1500m | *S.ae* | 4.921 | 1.223 | 16.378 | 4.025 | 0.005 |
|  | 1000m - 2000m | *S.ae* | 8.971 | 1.225 | 16.469 | 7.326 | 0.000 |
|  | 1500m - 2000m | *S.ae* | 4.050 | 1.218 | 16.108 | 3.326 | 0.020 |
|  | 500m - 1000m | *S.ch* | -1.623 | 1.234 | 17.017 | -1.315 | 0.567 |
|  | 500m - 1500m | *S.ch* | 1.187 | 1.255 | 18.153 | 0.946 | 0.781 |
|  | 500m - 2000m | *S.ch* | 4.535 | 1.250 | 17.886 | 3.627 | 0.010 |
|  | 1000m - 1500m | *S.ch* | 2.810 | 1.210 | 15.741 | 2.321 | 0.135 |
|  | 1000m - 2000m | *S.ch* | 6.158 | 1.215 | 15.974 | 5.067 | 0.001 |
|  | 1500m - 2000m | *S.ch* | 3.348 | 1.216 | 16.017 | 2.753 | 0.061 |

**Table S3** (cont’d)

| **Trait** | **contrast** | **Species** | **Estimate** | **SE** | **df** | **t ratio** | **P-value** |
| --- | --- | --- | --- | --- | --- | --- | --- |
| **Fig. 5h**  2019 SLA | 500m - 1000m | *S.ae* | -0.525 | 0.746 | 14.452 | -0.703 | 0.894 |
|  | 500m - 1500m | *S.ae* | 4.728 | 0.735 | 13.583 | 6.437 | 0.000 |
|  | 500m - 2000m | *S.ae* | 5.658 | 0.738 | 13.816 | 7.666 | 0.000 |
|  | 1000m - 1500m | *S.ae* | 5.253 | 0.750 | 14.730 | 7.007 | 0.000 |
|  | 1000m - 2000m | *S.ae* | 6.183 | 0.755 | 15.096 | 8.189 | 0.000 |
|  | 1500m - 2000m | *S.ae* | 0.930 | 0.734 | 13.523 | 1.267 | 0.597 |
|  | 500m - 1000m | *S.ch* | 1.301 | 0.746 | 14.400 | 1.743 | 0.338 |
|  | 500m - 1500m | *S.ch* | 3.605 | 0.739 | 13.834 | 4.880 | 0.001 |
|  | 500m - 2000m | *S.ch* | 4.186 | 0.747 | 14.407 | 5.602 | 0.000 |
|  | 1000m - 1500m | *S.ch* | 2.304 | 0.754 | 15.007 | 3.055 | 0.036 |
|  | 1000m - 2000m | *S.ch* | 2.885 | 0.765 | 15.752 | 3.773 | 0.008 |
|  | 1500m - 2000m | *S.ch* | 0.581 | 0.746 | 14.311 | 0.779 | 0.863 |
| **Fig. 5i**  2017 Total photosynthetic performance | 500m - 1000m | *S.ae* | -3.580 | 1.327 | 8.241 | -2.698 | 0.1000 |
|  | 500m - 1500m | *S.ae* | -4.358 | 0.438 | 8.923 | -9.942 | 0.0000 |
|  | 500m - 2000m | *S.ae* | -5.694 | 1.269 | 7.595 | -4.485 | 0.0099 |
|  | 1000m - 1500m | *S.ae* | -0.778 | 1.129 | 5.814 | -0.689 | 0.8976 |
|  | 1000m - 2000m | *S.ae* | -2.113 | 0.637 | 9.103 | -3.318 | 0.0368 |
|  | 1500m - 2000m | *S.ae* | -1.336 | 1.114 | 5.600 | -1.199 | 0.6504 |
|  | 500m - 1000m | *S.ch* | 0.003 | 1.313 | 7.970 | 0.002 | 1.0000 |
|  | 500m - 1500m | *S.ch* | -0.360 | 0.423 | 8.183 | -0.851 | 0.8293 |
|  | 500m - 2000m | *S.ch* | 0.008 | 1.266 | 7.533 | 0.006 | 1.0000 |
|  | 1000m - 1500m | *S.ch* | -0.363 | 1.118 | 5.631 | -0.325 | 0.9869 |
|  | 1000m - 2000m | *S.ch* | 0.005 | 0.620 | 8.385 | 0.008 | 1.0000 |
|  | 1500m - 2000m | *S.ch* | 0.368 | 1.115 | 5.626 | 0.330 | 0.9863 |
| **Fig. 5j**  2019 Chlorophyll | 500m - 1000m | *S.ae* | -7.912 | 1.378 | 18.885 | -5.741 | 0.000 |
|  | 500m - 1500m | *S.ae* | -6.477 | 1.309 | 15.663 | -4.947 | 0.001 |
|  | 500m - 2000m | *S.ae* | 3.007 | 1.299 | 15.154 | 2.316 | 0.138 |
|  | 1000m - 1500m | *S.ae* | 1.435 | 1.314 | 15.919 | 1.093 | 0.699 |
|  | 1000m - 2000m | *S.ae* | 10.920 | 1.379 | 18.923 | 7.916 | 0.000 |
|  | 1500m - 2000m | *S.ae* | 9.484 | 1.292 | 14.908 | 7.341 | 0.000 |
|  | 500m - 1000m | *S.ch* | -3.995 | 1.391 | 19.306 | -2.872 | 0.044 |
|  | 500m - 1500m | *S.ch* | 3.518 | 1.325 | 16.249 | 2.655 | 0.073 |
|  | 500m - 2000m | *S.ch* | 8.343 | 1.323 | 16.079 | 6.308 | 0.000 |
|  | 1000m - 1500m | *S.ch* | 7.513 | 1.324 | 16.272 | 5.674 | 0.000 |
|  | 1000m - 2000m | *S.ch* | 12.338 | 1.410 | 20.209 | 8.752 | 0.000 |
|  | 1500m - 2000m | *S.ch* | 4.825 | 1.323 | 16.121 | 3.648 | 0.010 |

**Table S3** (cont’d)

| **Trait** | **contrast** | **Species** | **Estimate** | **SE** | **df** | **t ratio** | **P-value** |
| --- | --- | --- | --- | --- | --- | --- | --- |
| **Fig. 5k**  2019 Flavonols | 500m - 1000m | *S.ae* | -0.119 | 0.048 | 19.341 | -2.469 | 0.097 |
|  | 500m - 1500m | *S.ae* | -0.213 | 0.049 | 20.575 | -4.329 | 0.002 |
|  | 500m - 2000m | *S.ae* | -0.413 | 0.046 | 16.775 | -8.908 | 0.000 |
|  | 1000m - 1500m | *S.ae* | -0.094 | 0.046 | 16.034 | -2.052 | 0.211 |
|  | 1000m - 2000m | *S.ae* | -0.294 | 0.049 | 20.607 | -5.985 | 0.000 |
|  | 1500m - 2000m | *S.ae* | -0.200 | 0.048 | 18.874 | -4.183 | 0.003 |
|  | 500m - 1000m | *S.ch* | 0.080 | 0.049 | 19.421 | 1.652 | 0.374 |
|  | 500m - 1500m | *S.ch* | 0.047 | 0.050 | 21.706 | 0.938 | 0.785 |
|  | 500m - 2000m | *S.ch* | 0.273 | 0.048 | 18.002 | 5.734 | 0.000 |
|  | 1000m - 1500m | *S.ch* | -0.033 | 0.046 | 16.224 | -0.719 | 0.888 |
|  | 1000m - 2000m | *S.ch* | 0.192 | 0.050 | 22.084 | 3.813 | 0.005 |
|  | 1500m - 2000m | *S.ch* | 0.225 | 0.050 | 20.980 | 4.538 | 0.001 |

**Fig. S1** The number of differentially expressed genes with distance from each species’ home site. Analyses performed in limma/voom represented as unfilled circles and dashed lines, and DESeq2 as solid circles and lines.

**Table S4** Significantly enriched Gene Ontology terms representing differentially expressed genes between the highest and lowest transplant site in *S. aethnensis*

| **GO.ID** | **Term** | **Annotated** | **Significant** | **p-value** |
| --- | --- | --- | --- | --- |
| GO:0072657 | Protein localization to membrane | 25 | 13 | 0.02 |
| GO:0009226 | nucleotide-sugar biosynthetic proces | 8 | 6 | 0.015 |
| GO:0046364 | monosaccharide biosynthetic process | 6 | 5 | 0.015 |
| GO:0016117 | carotenoid biosynthetic process | 10 | 8 | 0.002 |
| GO:0046500 | S-adenosylmethionine metabolic process | 6 | 5 | 0.014 |
| GO:0042744 | hydrogen peroxide catabolic process | 8 | 6 | 0.015 |
| GO:0006555 | methionine metabolic process | 8 | 6 | 0.015 |
| GO:0015706 | nitrate transport | 7 | 6 | 0.005 |
| 0000027 | ribosomal large subunit assembly | 7 | 5 | 0.037 |
| 0009624 | response to nematode | 5 | 4 | 0.038 |
| 0042743 | hydrogen peroxide metabolic process | 11 | 9 | 0.009 |
| 0017004 | cytochrome complex assembly | 11 | 7 | 0.03 |

**Table S5** Significantly enriched Gene Ontology terms representing differentially expressed genes between the highest and lowest transplant site in *S. chrysanthemifolius*

| **GO.ID** | **Term** | **Annotated** | **Significant** | **p-value** |
| --- | --- | --- | --- | --- |
| GO: 0009226 | nucleotide-sugar biosynthetic process | 8 | 8 | 0.005 |
| GO: 0019321 | pentose metabolic process | 6 | 6 | 0.01 |
| GO: 0042744 | hydrogen peroxide catabolic process | 8 | 7 | 0.04 |
| GO: 0007018 | microtubule-based movement | 10 | 9 | 0.01 |
| GO: 0045490 | pectin catabolic process | 6 | 6 | 0.01 |
| GO: 0000027 | ribosomal large subunit assembly | 7 | 7 | 0.009 |
| GO: 0042737 | drug catabolic process | 28 | 24 | 0.0001 |
| GO: 0006412 | translation | 127 | 76 | 0.03 |
| GO: 0009895 | negative regulation of catabolic process | 9 | 8 | 0.024 |
| GO: 0010928 | regulation of auxin mediated signaling | 5 | 5 | 0.036 |
| GO: 0033500 | carbohydrate homeostasis | 5 | 5 | 0.036 |
| GO: 0098754 | detoxification | 6 | 6 | 0.018 |
| GO: 1903530 | regulation of secretion by cell | 5 | 5 | 0.036 |
| GO: 0006518 | peptide metabolic process | 134 | 82 | 0.013 |

**Table S6** Reconstructed WGCNA network modules. ‘No. genes’ represents the number of genes for each module. The correlation (Pearson’s r) between the module eigengene and elevation is presented in the top cell for each module with cell colour representing the direction of the correlation (red for positive and green for negative) and shading indicating the strength of the correlation. Adjusted p-value testing the significance of the correlation with elevation are presented in the lower cell for each module.

**Methods S1**

*Environmental conditions of the growth cabinet for the common garden experiment*
350μmol m^-2^ s^-1^ light intensity, 25/20°C±3°C day/night temperature, 65-70% humidity and 14/10h photoperiod and 400μmol mol^-1^ ambient CO_2_ concentration.

*Calculations of physiological light defence in the common garden experiment*
Using output from the fluorometer, we quantified two mechanisms of physiological light defence: (1) the unregulated dissipation of heat as the quantum yield of non-regulated energy dissipation, calculated as $Y\left( NO \right)=\frac{1}{NPQ+1+qL(\frac{F_{m}}{F_{o}}-1)}$, where $qL$ is the parameter representing photochemical quenching and $NPQ=\frac{F_{m}-F_{m}^{'}}{F_{m}^{'}}$, where $F_{m}$ and $F_{m}^{'}$ represent the maximal fluorescence in a dark and light adapted state, respectively (Kramer et al. 2004). (2) To quantify the regulated dissipation of heat from leaves, we calculated the quantum yield of regulated energy dissipation as $Y\left( NPQ \right)=1-Y\left( II \right)-Y\left( NO \right)$, where $Y\left( II \right)$ is the efficiency of photosystem II (Kramer et al. 2004).

*Calculations of total photosynthetic performance in the field transplant*
Using the JIP test (Tsimilli-Michael and Strasser 2013), we calculated the total performance of photosystem I and II as: ${PI}_{total}=\frac{F_{V}}{F_{M}}\times\frac{V_{J}}{M_{0}}\times\frac{F_{V}}{F_{0}}\times\frac{1-V_{J}}{V_{J}}\times\frac{1-V_{I}}{V_{I}-V_{J}}$, where F_0_ is the minimal fluorescence intensity, F_M_ the maximal fluorescence, F_V_ the maximal variable fluorescence (F_V_ = F_M_ – F_0_), M_0_ the approximated initial slope of fluorescence change (normalised on F_V_), and V_J_ and V_I_ the relative variable fluorescence levels recorded at 2ms and 30ms, respectively (also normalised on F_V_).

*Estimation of D-matrix to visualise differences in multivariate phenotype*
We calculated **D** using $\mathbf{D}=\frac{\mathrm{MS}_{H}-\mathrm{MS}_{E}}{nf}$, where the mean square matrices (MS) represent the differences in mean multivariate phenotype among species and elevations (MS_H_) and the residual error (MS_E_). MS are calculated by extracting the sums-of-squares and cross-product matrices (SSCP) from the MANOVA, and dividing by the appropriate degrees of freedom. In our design, we compared both species across the four elevations (MS_H_ *df* = 7), and in the MANVOA we designated genotypes (within species) measured at all four elevations as the error term (MS_E_ *df* = 298 in 2017 and *df =* 176 in 2019). This tests whether differences among species and elevations are greater than among genotypes measured at all elevation. The denominator (*nf*) represents the average number of individuals measured for each genotype across elevations, which is calculated according to equation 9 in Martin et al. (2008). Analogous to principal components scores, we then calculated the scores of the first two axes of **D**, which provided a visualisation for how multivariate phenotype changed across elevation for both species.

*Transcriptome assembly*
Raw reads were processed in *TrimGalore* v0.6 to remove low quality bases and adaptors (Phred quality cut-off score = 20). For each species, trimmed reads from all samples were combined and a reference transcriptome was *de novo* assembled using *Trinity* v2.8.4 (Haas et al. 2013). Each transcriptome was filtered using *EvidentialGene* (<http://arthropods.eugenes.org/EvidentialGene/>) and contaminants were removed using *MCSC Decontamination* (Lafond-Lapalme et al. 2017), with the filter set to ‘Viridiplantae’. Each transcriptome was then filtered to contain only orthologous transcripts identified using *Orthofinder* v2.3.3 with default parameters (Emms and Kelly 2019). The final orthologous transcriptomes each contained 23,622 transcripts with an N50 of 1,509 and 1,515 for *S. aethnensis* and *S. chrysanthemifolius,* respectively.

*Annotation of differentially expressed genes*
Reference transcriptomes were annotated using *Trinotate* v3.2.1 (Bryant et al. 2017). Predicted amino-acid sequences were generated using *TransDecoder* v5.5 (<https://transdecoder.github.io>) and protein sequences were blasted against the Uniprot database. Annotation of the transcriptomes resulted in 7,579 unique GO (Gene Ontology) terms for 14,701 transcripts (mean of 7.2 GO terms/transcript).

*Weighted network construction of differentially expressed genes*
Transformed expression data were split by species and analysed using *WGCNA* v1.51 (Langfelder and Horvath 2008). For each species, a similarity matrix between all genes was first calculated using Pearson’s correlation, and then adjacencies between genes were estimated using a soft-threshold power of β=6. A Topological Overlap Matrix (TOM) was then estimated by identifying genes that were similar in terms of shared neighbours. Each TOM was scaled such that the 95^th^ percentiles were equivalent and a consensus TOM was calculated from the minimum topological overlap. Modules of highly coexpressed genes were identified using hierarchical clustering and refined with a dynamic tree cut so that similar modules were collapsed into one (Langfelder and Horvath 2008).

*References*

Bryant, D. M., K. Johnson, T. DiTommaso, T. Tickle, M. B. Couger, D. Payzin-Dogru, T. J. Lee, N. D. Leigh, T. H. Kuo, F. G. Davis, J. Bateman, S. Bryant, A. R. Guzikowski, S. L. Tsai, S. Coyne, W. W. Ye, R. M. Freeman, L. Peshkin, C. J. Tabin, A. Regev, B. J. Haas, and J. L. Whited. 2017. A Tissue-Mapped Axolotl De Novo Transcriptome Enables Identification of Limb Regeneration Factors. Cell Reports 18:762-776.

Emms, D. M. and S. Kelly. 2019. OrthoFinder: phylogenetic orthology inference for comparative genomics. bioRxiv:466201.

Haas, B. J., A. Papanicolaou, M. Yassour, M. Grabherr, P. D. Blood, J. Bowden, M. B. Couger, D. Eccles, B. Li, M. Lieber, M. D. MacManes, M. Ott, J. Orvis, N. Pochet, F. Strozzi, N. Weeks, R. Westerman, T. William, C. N. Dewey, R. Henschel, R. D. Leduc, N. Friedman, and A. Regev. 2013. De novo transcript sequence reconstruction from RNA-seq using the Trinity platform for reference generation and analysis. Nature Protocols 8:1494-1512.

Kramer, D. M., G. Johnson, O. Kiirats, and G. E. Edwards. 2004. New fluorescence parameters for the determination of QA redox state and excitation energy fluxes. Photosynthesis Res. 79:209-218.

Lafond-Lapalme, J., M. O. Duceppe, S. R. Wang, P. Moffett, and B. Mimee. 2017. A new method for decontamination of de novo transcriptomes using a hierarchical clustering algorithm. Bioinformatics 33:1293-1300.

Langfelder, P. and S. Horvath. 2008. WGCNA: an R package for weighted correlation network analysis. BMC Bioinformatics 9:559.

Martin, G., E. Chapuis, and J. Goudet. 2008. Multivariate Q_st_–F_st_ Comparisons: A Neutrality Test for the Evolution of the G Matrix in Structured Populations. Genetics 180:2135-2149.

Tsimilli-Michael, M. and R. J. Strasser. 2013. Biophysical Phenomics: Evaluation of the Impact of Mycorrhization with *Piriformospora indica*. Pp. 173-190 *in* A. Varma, G. Kost, and R. Oelmüller, eds. *Piriformospora indica*: Sebacinales and their biotechnological applications. Springer, Berlin, Germany.
